# Deletions Rate-Limit Breast and Ovarian Cancer Initiation

**DOI:** 10.1101/2024.10.17.618945

**Authors:** Kathleen E. Houlahan, Mahad Bihie, Julián Grandvallet Contreras, Daniel J. Fulop, Gonzalo Lopez, Hsin-Hsiung Huang, Peter Van Loo, Christina Curtis, Paul C. Boutros, Kuan-lin Huang

## Abstract

Optimizing prevention and early detection of cancer requires understanding the number, types and timing of driver mutations. To quantify this, we exploited the elevated cancer incidence and mutation rates in germline *BRCA1* and *BRCA2 (gBRCA1/2)* carriers. Using novel statistical models, we identify genomic deletions as the likely rate-limiting mutational processes, with 1-3 deletions required to initiate breast and ovarian tumors. *gBRCA1/2*-driven hereditary and sporadic tumors undergo convergent evolution to develop a similar set of driver deletions, and deletions explain the elevated cancer risk of *gBRCA1/2*-carriers. Orthogonal mutation timing analysis identifies deletions of chromosome 17 and 13q as early, recurrent events. Single-cell analyses confirmed deletion rate differences in *gBRCA1/2* vs. non-carrier tumors as well as cells engineered to harbor *gBRCA1/2*. The centrality of deletion-associated chromosomal instability to tumorigenesis shapes interpretation of the somatic evolution of non-malignant tissue and guides strategies for precision prevention and early detection.

## INTRODUCTION

Cancer arises through the accumulation of somatic mutations^1,2^, but the specific mutational processes and number of molecular alterations required for tumor initiation remain unclear. Several studies have identified accumulations of cancer-associated mutations in normal tissues^12,13^. Further, large numbers of pre-malignant lesions are diagnosed each year and must be risk-stratified to identify their likelihood of progression to aggressive cancers. Thus, understanding the fundamental rate-limiting steps required for tumorigenesis remains a critical unresolved problem in cancer biology.

Both theoretical and observational methods have been proposed to address this fundamental question^3–11^. Theoretical models that relate the number of mutations in driver genes to the distribution of age-at-diagnosis suggest 2-10 drivers are required, based on differing assumptions on drivers, pathways, and environmental influences^3–6^. Observational approaches have been developed to distinguish somatic driver mutations from background passenger alterations, including recurrent point mutations^7^, positively-selected point mutations^8^, aberrant methylation^9^, genomic rearrangements^10^ and copy number aberrations (CNAs)^11^.

Seminal studies by Nordling^14^ and Armitage and Doll^15^ demonstrated that the number of aberrations required for tumorigenesis could be inferred from the relationship between age and cancer incidence. Knudson famously compared the incidence curves for familial and sporadic retinoblastoma, resulting in the “two-hit” hypothesis for *RB1*^16^, along with the more recent discovery of “one-hit” early-onset retinoblastoma^17^. Tomasetti and colleagues extended this framework to estimate driver count from the differential mutation and incidence rates between two cancer sub-groups^6^. They leveraged differing single nucleotide variant (SNV) mutation rates estimated from DNA sequencing and epidemiological incidence rates to estimate the minimum number of sequential SNVs required to initiate a cancer. Comparing smokers *vs.* non-smokers in lung cancer and microsatellite instable (MSI) *vs*. microsatellite stable (MSS) colorectal cancer, they identified a minimum of three sequential SNVs were required for both diseases.

*BRCA1* and *BRCA2* germline variant carriers are strongly predisposed to cancers in multiple organs, including prostate, pancreatic, breast and ovarian cancer^18,19,20^. Similar to lung tumors in smokers and MSI colorectal tumors—tumors in *BRCA1* and *BRCA2* carriers show elevated mutation rates^21–25^, at least in part a result of their homologous recombination deficiency (HRD)^26^. Women with pathogenic germline variants in these genes are often intensively screened, and pre-malignant lesions are common in both the breast and ovaries^27–30^. To understand the initiation of these cancers that are not driven by point mutations, we developed a quantitative driver-estimation framework based on statistical formulations by Tomasetti *et al.*^6^ to estimate the minimum number of SNV, short insertions and deletion (INDEL) and CNA drivers. We applied this framework to 1256 breast and ovarian sporadic and BRCA-driven cancer cases, identifying deletions as the rate-limiting mutational process leading to breast and ovarian tumorigenesis – contrary to the emphasis placed on point mutations in these tumors. We confirm these observations using both driver prioritization^31,32^ and reconstruction of clonal evolution^33^. Taken together, we provide a new framework for integrating theoretical and empirical approaches to driver mutation requirements across all types of driver mutation. We demonstrate its utility for HRD-cancers, informing early-detection and risk-stratification approaches in breast and ovarian cancer.

## RESULTS

### *BRCA1* and *BRCA2* carriers showed varied rates of somatic alterations

While the role of *BRCA1* and *BRCA2* in homologous recombination has been well characterized, the consequence of variants in these two genes is lineage dependent^20^. We focus here on two canonical *BRCA*-associated cancers, those arising in the breast and ovary. Leveraging data from TCGA Pan-Cancer Atlas^34,35^, we interrogated the somatic mutation landscape of 758 breast and 498 ovarian tumors, a subset of which harboured pathogenic germline variants in *BRCA1* (n_breast_ = 20; n_ovarian_ = 31) or *BRCA2* (n_breast_ = 12; n_ovarian_ = 22) as evaluated by CharGer and the PanCanAtlas germline study (Characterization of Germline Variants; sensitivity for detecting pathogenic variants = 88%; false-positive rate = 4.9%)^35^. We only consider individuals of European descent to avoid confounding by ancestry-biased somatic mutations^36^. There were no significant differences in tumor purity or ploidy between carriers and non-carriers (**Supplementary Figure 1**), aside from *BRCA2* loss-of-function ovarian samples, which had significantly lower tumor purity than non-carrier samples. The lower purity suggests our mutation estimates for this subgroup are conservative (because of reduced coverage) and thus serve as a lower bound.

We first compared the burden of diverse genomic alterations, including SNVs, CNAs, and INDELs, in *BRCA1* and *BRCA2* carriers *versus* non-carriers. As expected^21–24^, somatic SNV mutation counts, normalized by age at diagnosis and adjusted for stage, grade and subtype, were elevated in breast and ovarian tumors arising in individuals carrying pathogenic germline variants in *BRCA1* (β_breast_ = 0.30 (95% Confidence Intervals(CI): 0.11-0.49); FDR_breast_ = 1.9×10^-2^; β_ovarian_ = 0.23 (95% CI: 0.12-0.34); FDR_ovarian_ = 3.8×10^-3^; linear regression) or *BRCA2* (β_breast_ = 0.24 (95% CI: -0.01-0.50); FDR_breast_ = 0.19; β_ovarian_ = 0.27 (95% CI: 0.13-0.41); FDR_ovarian_ = 5.7×10^-3^; **Figure 1a-c**).

**Figure 1:**
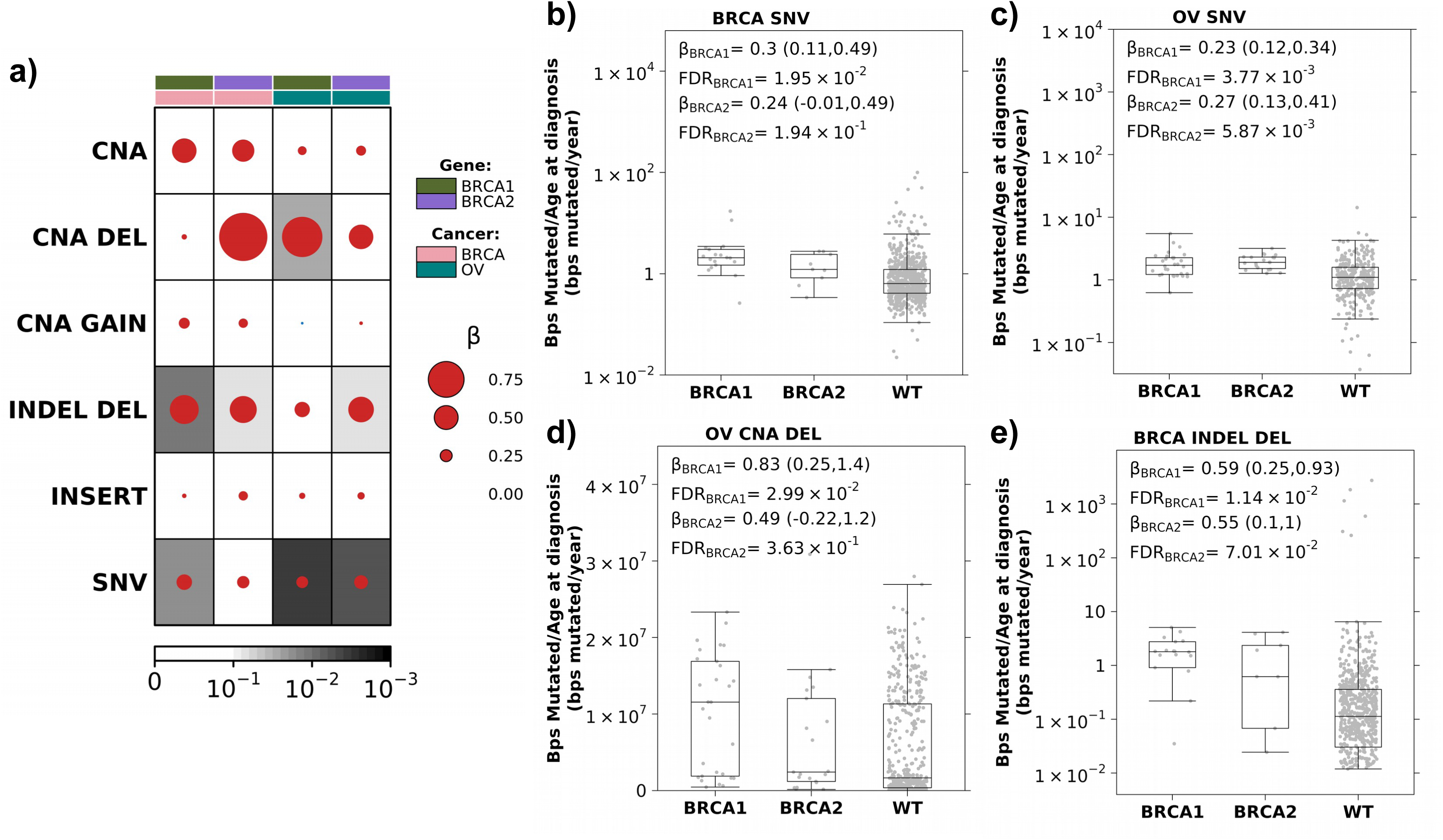
Increased mutation burden in *BRCA1* and *BRCA2* carriers. **a)** Breast and ovarian tumors in *BRCA1* and *BRCA2* carriers show an increased mutation rate (bp/year) compared to non-carrier (WT) individuals. Dot size and colour reflect β magnitude and direction from linear regression correcting for stage, grade and subtype. Background shading indicates FDR. Covariate along the top indicates gene and cancer type. **b-c)** Boxplots show increased coding SNV mutation rate (bp/year) in breast **(b)** and ovarian **(c)** tumors in *BRCA1* and *BRCA2* carriers compared to WT. Boxplots represent median, 0.25 and 0.75 quantiles with whiskers are 1.5x interquartile range. β and FDR from linear regression. **d)** *BRCA1* and *BRCA2* carriers show an increased CNA deletion rate compared to WT in ovarian tumors. **e)** *BRCA1* and *BRCA2* carriers show an increased small deletion rate compared to WT in breast tumors. NV: single nucleotide variants; CNA: copy number aberrations; DEL: deletions; INDEL: small insertion and deletions

Because CNAs also accumulate with age^37^, we considered the total number of base pairs involved in CNAs, normalized by age at diagnosis. After controlling for stage, grade and subtype, there was no significant difference in CNA burden between carriers and non-carriers (FDR > 0.1; **Figure 1a**). Considering gains (GAIN) and deletions (DEL) separately, ovarian tumors in *BRCA1* carriers had a significantly higher CNA deletion burden than non-carriers (β = 0.83 (95% CI: 0.25-1.4); FDR = 3.0×10^-2^; **Figure 1a&d**) while no significant difference in CNA gains was observed between carriers and non-carriers (FDR ≥ 0.64; **Supplementary Figure 2**).

Next, we considered INDELs (<1kbp) based on consensus calling by the TCGA PanCanAtlas MC3 project^34^. We calculated INDEL burden as the number of base pairs involved in an INDEL normalized by age at diagnosis and similarly quantified a significant increase in INDEL deletions in *BRCA1* (β_breast_= 0.59 (95% CI: 0.25-0.93); FDR_breast_ = 1.14×10^-2^; β_ovarian_ = 0.30 (95% CI: -0.02-0.62); FDR_ovarian_ = 0.19) and *BRCA2* (β_breast_ = 0.55 (95% CI: 0.10-1.0); FDR_breast_ = 6.5×10^-2^; β_ovarian_ = 0.52 (95% CI: 0.11-0.92); FDR_ovarian_ = 6.4×10^-2^) carriers (**Figure 1a&e**). No significant increased burden of INDEL insertions was observed (FC ≤ 0.18; FDR ≥ 0.31; **Supplementary Figure 2**). Thus, breast and ovarian tumors in *BRCA1* and *BRCA2* carriers recapitulate the elevated burden of diverse genomic alterations expected of this high-risk group.

### Accounting for increased incidence ratio of *BRCA* carriers

Expanding upon the insights of Nordling, Armitage and Doll^14,15^, Tomasetti *et al.* showed cancer incidence is a function of the number of rate-limiting mutational events required for initiation of a cancer subgroup and the average mutation rate of that mutation type in that cancer subgroup^6^. This framework assumes a constant mutation rate over evolutionary time. For example, a cancer subgroup that has a mutation burden twice that of another subgroup will have an incidence rate 2^n^ that of the second subgroup, where n represents the number of rate-limiting mutational events required for tumorigenesis (**Methods**). *BRCA1* and *BRCA2* are key components of the DNA damage repair pathway; their pathogenic germline variants lead to deficient homologous-directed repair. We hypothesize that the elevated incidence rates in *BRCA1* and *BRCA2* carriers can, in part, be explained by their increased mutational burden. We therefore estimated the number of driver events required to initiate breast and ovarian cancer by comparing *BRCA1* and *BRCA2* germline variant carriers to non-carriers (**Methods**). By expanding the framework to consider multiple somatic alteration types, we can quantify the number of each alteration type required to converge on the observed incidence rates and identify the rate-limiting mutational processes. Identifying the rate-limiting mutational processes shapes how we understand tumor evolution.

We first reviewed the epidemiology literature to obtain standard incidence ratios (SIRs) of breast and ovarian cancer in *BRCA1* and *BRCA2* carriers compared to matched control populations (**Supplementary Table 1**). The best powered study, including 6,036 *BRCA1* and *BRCA2* female carriers, revealed SIRs of: SIR_breast|*BRCA1*_ (95% CI): 16.6 (14.7-18.7), SIR_breast*BRCA2*_: 12.9 (11.1-15.1), SIR_ovarian|*BRCA1*_: 49.6 (40.0-61.5), SIR_ovarian|*BRCA2*_: 13.7 (9.1-20.7)^38^.

Next, we conducted a bootstrapped estimation (n=10,000) of the average increase in SNV mutation burden (*i.e.* protein-altering SNVs counts normalized by age at diagnosis) for *BRCA1* and *BRCA2* carriers compared to clinically-matched non-carrier (WT) individuals with breast and ovarian cancer. We identified the expected increase in normalized SNV counts in carriers (median FC_breast|*BRCA1*_ *_vs._* _WT_ (95 CI%) = 3.57 (2.17-5.19); median FC_breast|*BRCA2*_ *_vs._* _WT_ = 2.21 (1.71-4.06); median FC_ovarian|*BRCA1*_ *_vs._* _WT_ = 1.60 (1.13-2.16); median FC_ovarian|*BRCA2*_ *_vs._* _WT_ = 1.82 (1.31-2.61); **Supplementary Figure 3a**). Finally, we compared the estimated mutation burden ratios to observed SIRs for each pair of germline risk-gene and cancer type (**Figure 2&3**).

**Figure 2:**
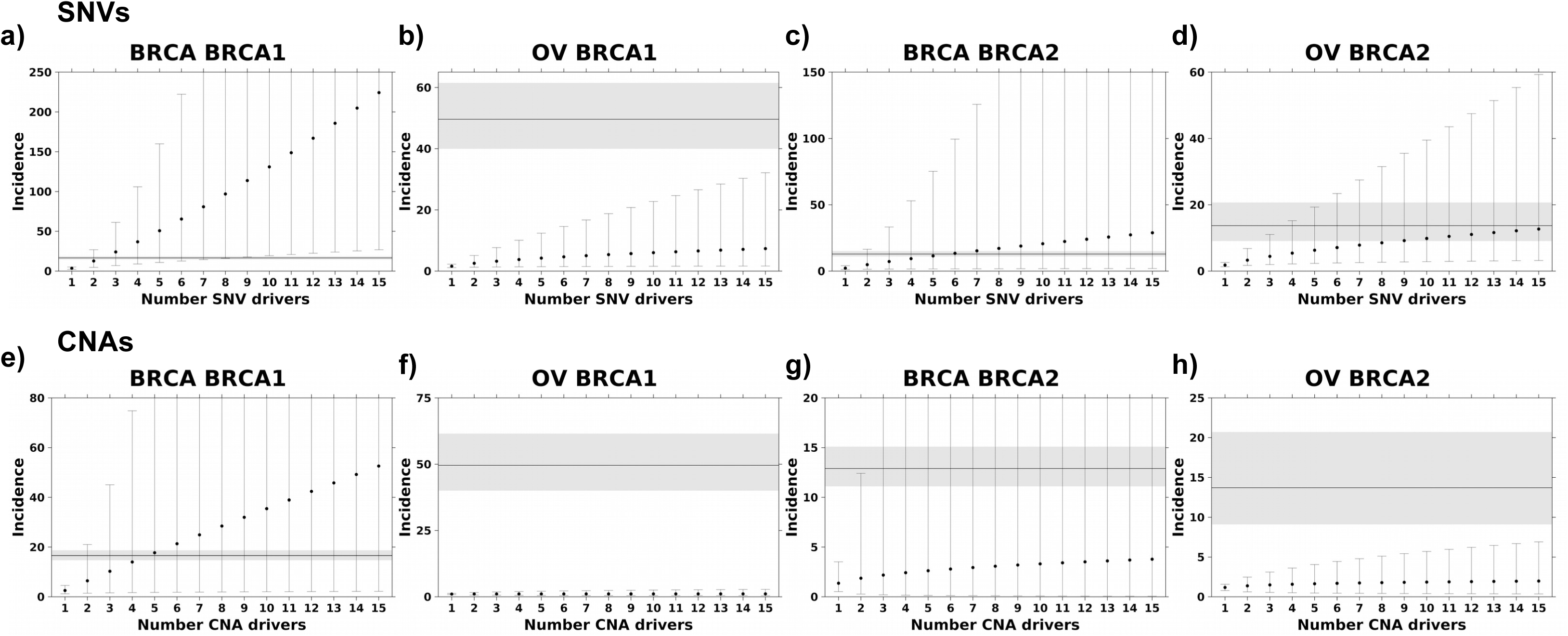
SNV and CNA mutation burden alone cannot explain breast and ovarian cancer incidence. Estimated incidence rates based on SNV **(a-d)** and CNA **(e-h)** mutational burden with the increasing number of drivers cannot explain the observed incidence ratio of breast and ovarian cancer in *BRCA1* and *BRCA2* carriers. Points represent estimated incidence ratios based on the number of driver events along the x-axis for breast cancer in *BRCA1* **(a&e)** and *BRCA2* **(b&f)** carriers and ovarian cancer in *BRCA1* **(c&g)** and *BRCA2* **(d&h)** carriers. Error bars indicate 95% confidence intervals. The horizontal line indicates observed incidence rate, and grey shading indicates 95% confidence intervals.

**Figure 3:**
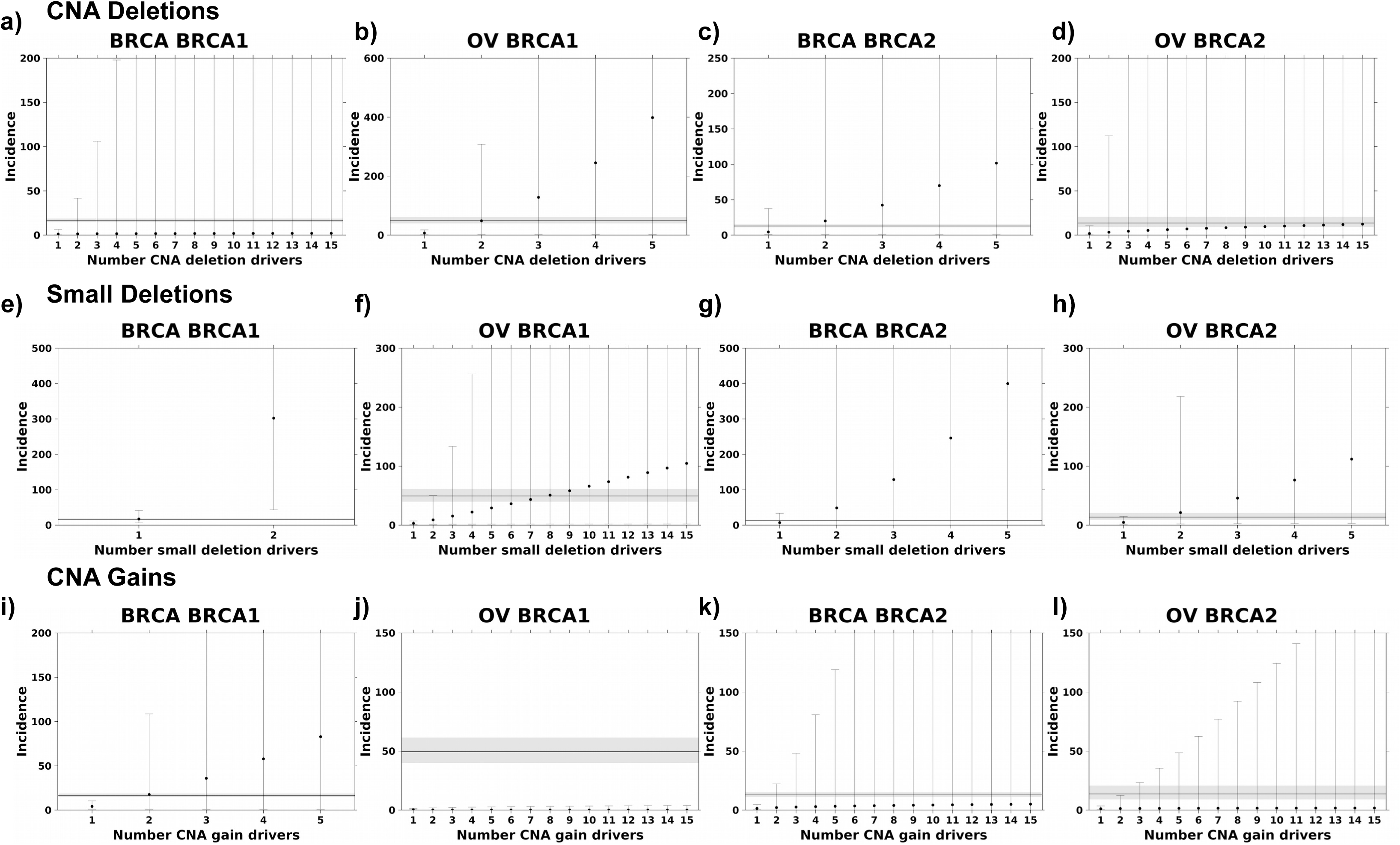
Deletions are the likely rate-limiting mutational process in *BRCA1* and *BRCA2* carriers. **a-d)** Incidence estimates based on deletion rates converge on the observed incidence rate. Deletion burden calculated as the number of base pairs altered normalized by age at diagnosis. **e-h)** Incidence estimates based on small deletions. **i-l)** Incidence estimates based on gain rates. Segplots compare estimated incidence rate to observed incidence rate as described in Figure 2.

For breast cancer, *BRCA1* germline variant carriers had 3.57-fold more protein-altering SNVs than *BRCA1* WT patients. Thus, if breast cancer initiation in these individuals only required two driver SNVs, the estimated incidence ratio would be 3.57^2^=12.75. If three coding SNV drivers are required, the estimated incidence ratio would be 3.57 x 3.57 x 3.57^1/2^ = 24.08 (see Methods). The observed SIR for breast cancer *BRCA1* carriers is 16.6; therefore, the framework estimated a minimum of two SNVs are required to initiate breast cancer in *BRCA1* carriers (D = 0.30; Kolmogorov–Smirnov test; **Figure 2a**). The framework estimates five to ten SNV drivers are required to initiate breast and ovarian cancer in *BRCA2* carriers (D_breast_(5 drivers) = 0.17; D_ovarian_(10) = 0.20; Kolmogorov–Smirnov test; **Figure 2c-d**), however, it fails to converge on the ovarian SIR in *BRCA1* carriers (Kolmogorov–Smirnov D did not reach minimum; **Figure 2b**). These data suggest the SNV mutation burden does not fully capture the mechanisms of germline *BRCA1* and *BRCA2* deficiency and SNVs are likely not the rate-limiting mutational process leading to tumorigenesis in *BRCA1* and *BRCA2* carriers.

Next, we explored other mutational processes that may explain the increase in cancer incidence in *BRCA1* and *BRCA2* carriers. Given that ovarian tumors are C-class tumors characterized by high CNA rates^11^, we evaluated if elevated CNA burden, *i.e.* CNA bp/year, could explain breast and ovarian cancer incidence in *BRCA1* and *BRCA2* carriers. Compared to WT, *BRCA1* and *BRCA2* carriers had modest increases in CNA burden (median FC_breast|*BRCA1*_ *_vs._* _WT_ (95 CI%) = 1.26 (1.00-1.54); median FC_breast|*BRCA2*_ *_vs._* _WT_ = 1.06 (0.83-1.76); median FC_ovarian|*BRCA1*_ *_vs._* _WT_ = 1.19 (1.06-1.37); median FC_ovarian|*BRCA2*_ *_vs._* _WT_ = 1.15 (1.03-1.34); **Supplementary Figure 3b**). Our framework estimates four CNA drivers are required to initiate breast cancer in *BRCA1* carriers (D = 0.17; Kolmogorov–Smirnov test; **Figure 2e**), however, the framework failed to reach the observed SIR in the other three cancer-gene pairs (Kolmogorov–Smirnov D did reach minimum; **Figure 2f-h**). Driver estimates were similar calculating CNA burden as the number of CNA segments normalized by age at diagnosis (**Supplementary Figure 4a-d**). A CNA driver could be a single event deleting or amplifying multiple genes in close proximity, *e.g. RB1* and *BRCA2* on chromosome 13, or a single event capturing a single driver gene and many passenger genes.

Pathogenic *BRCA1* and *BRCA2* variants induce homologous repair deficiency (HRD), which has been associated with increased rates of genomic deletions^10,22,39^. Thus, we next considered deletions and gains separately. Considering only deletions and calculating mutation burden based on the number of deletions or base pairs altered, the ratio of CNA mutation rates converged on the observed SIR in both breast and ovarian cancer (**Figure 3a-d; Supplementary Figure 4e-h**). *BRCA1* and *BRCA2* carriers had increased deletion mutation rates (bp/year) compared to non-carrier individuals (median FC_breast|*BRCA1*_ *_vs._* _WT_ (95 CI%) = 1.18 (0.28-6.46); median FC_breast|*BRCA2*_ *_vs._* _WT_ = 4.48 (0.83-37.6); median FC_ovarian|*BRCA1*_*_vs._*_WT_ = 6.97 (1.26-17.6); median FC_ovarian|*BRCA2*_ *_vs._* _WT_ = 1.80 (0.32-10.6); **Supplementary Figure 3c**). Modeling based on CNA deletions estimated two to four deletions might be required to initiate breast or ovarian cancer in *BRCA1* and *BRCA2* carriers (D_ovarian|*BRCA1*_(2) = 0.16; D_breast|*BRCA2*_(2) = 0.31; D_ovarian|*BRCA2*_(4) *=* 0.23). The framework did not converge on the SIR for breast cancer in *BRCA1* carriers suggesting CNA gains may account for the estimated four CNA drivers in this subgroup (**Figure 2e, 3a**).

Next, we considered small deletions (< 1kbp) and similarly observed an increased rate of small deletions (bp/year) in the *BRCA1* and *BRCA2* carriers compared to non-carrier individuals (median FC_breast|*BRCA1*_ *_vs._* _WT_ (95 CI%) = 17.4 (6.60-41.5); median FC_breast|*BRCA2*_ *_vs._* _WT_ = 6.98 (0.42-33.7); median FC_ovarian|*BRCA1*_ *_vs._* _WT_ = 3.00 (1.16-7.08); median FC_ovarian|*BRCA2*_ *_vs._* _WT_ = 4.62 (1.41-14.8); **Supplementary Figure 3d**). Based on small deletions, considering both number of deletions and base pairs altered, we estimate one to two small deletions are required for breast or ovarian tumorigenesis in *BRCA1* and *BRCA2* carriers (D_breast|*BRCA1*_(1) = 0.25; D_ovarian|*BRCA1*_(6) = 0.16; D_breast|*BRCA2*_(1) = 0.31; D_ovarian|*BRCA2*_(2) *=* 0.27; **Figure 3e-h; Supplementary Figure 4i-l**). In contrast, *BRCA1* and *BRCA2* carriers did not show a consistent increase in gain rates (median FC_breast|*BRCA1*_ *_vs._*_WT_ (95 CI%) = 4.19 (0.90-10.4); median FC_breast|*BRCA2*_ *_vs._* _WT_ = 1.46 (0.26-4.71); median FC_ovarian|*BRCA1*_ *_vs._* _WT_ = 0.37 (0.25-1.38); median FC_ovarian|*BRCA2*_ *_vs._* _WT_ = 1.14 (0.30-3.52); **Supplementary Figure 3e**). Modeling based on CNA gains estimated two drivers are required in *BRCA1*-driven breast cancer (D(2) = 0.16; **Figure 3i**), however, the framework failed to reach the observed SIR in the other three cancer-gene pairs (Kolmogorov–Smirnov D did reach minimum; **Figure 3j-l**). Driver estimates were similar calculating CNA gain burden as number of CNA gain segments normalized by age at diagnosis (**Supplementary Figure 4m-p**). Because of the large number of samples with no small insertions, the framework could not be applied to small insertions.

We next replicated this statistical analysis using the METABRIC cohort of 1,980 breast tumors39, which included 26 *BRCA1* and 31 *BRCA2* carriers (Methods). The framework failed to reach the observed SIR when considering total SNV and CNA burden (Supplementary Figure 5). Only when considering the number of genes lost did the model converge for both BRCA1- and BRCA2-driven breast cancer, aligned with the observation in TCGA. Our model predicted 4 and 9 gene drivers were required in *BRCA1*-(D(4) = 0.17) and *BRCA2*-(D(9) = 0.18) driven tumors, respectively, in METABRIC.

These data suggest deletions are likely the predominant rate-limiting mutational process leading to breast and ovarian tumorigenesis if tumours are initiated based on the least numbers of required drivers (**Figure 4a**). However, an assumption of our framework is that a mechanism by which *BRCA1* and *BRCA2* germline variants increase the risk of tumorigenesis is through suppression of DNA damage repair leading to increased mutational burden^21–25^. If *BRCA1* and *BRCA2* tumors have different driver type requirements than non-carrier tumors, comparison of alteration rates may be invalid. To evaluate the possibility, we compared CNA patterns of *BRCA1*, *BRCA2* and non-carrier tumors for both cancer types (**Methods**). Breast and ovarian *BRCA1* and *BRCA2* tumors have higher frequencies of CNAs but carriers had the same CNA patterns as non-carrier tumors (**Supplementary Figure 6**). For example, the most recurrent CNAs in breast cancer include chromosome 17p deletion (frequencies in *BRCA1*: 30.0%; *BRCA2*: 63.6%;, WT: 37.7%) and chromosome 8q gain (frequencies in *BRCA1*: 70.0%; *BRCA2*: 72.7%;, WT: 45.8%). The most recurrent CNAs in ovarian cancer include 8q gain (frequencies in *BRCA1*: 79.3%; *BRCA2*: 75.4%; WT: 62.7%). The similar CNA landscapes suggest *BRCA1*, *BRCA2* and non-carrier tumors require similar deletions and gains while acquiring these alterations at different rates. Both BRCA-driven hereditary and sporadic tumors evolve to converge on a similar tumorigenic state. Based on these results, we concluded our approach is justified.

**Figure 4:**
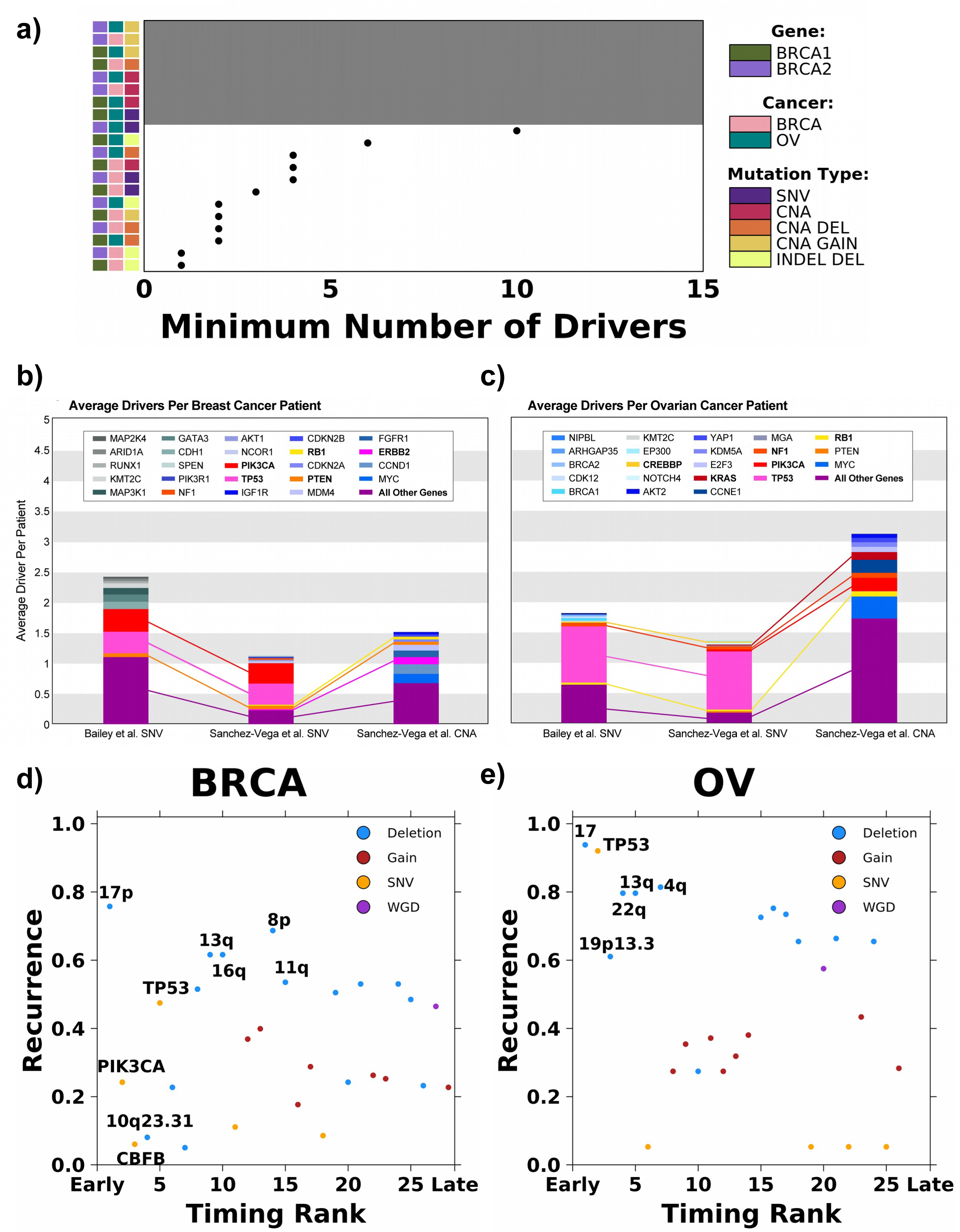
Empirical driver and timing prediction from sequencing studies. **a)** Summary of the minimum number of drivers estimated for each cancer type, carrier and mutation type modeled. Grey background indicates the model did not converge and covariate along the left indicates cancer type, carrier and mutation type modeled. **b-c)** The average number of drivers per patient observed for each gene from sequencing data in TCGA breast and ovarian cancers. The top 10 genes with the highest average driver count per patient are colored distinctively, and all other driver genes are grouped into a single category. Connections between stacked bars indicate the same genes in the top 10 across the driver prioritization method and variant type. Each stacked bar signifies the average total driver SNV or CNA found amongst patients with breast **(b)** and ovarian **(c)** cancers. **d-e)** Recurrence (y-axis) and relative timing (early *vs.* late; x-axis) for 28 and 26 drivers in breast **(d)** and ovarian **(e)** cancer, respectively. Drivers are colored based on the type of mutational event. The union of the top five most recurrent and earliest occurring drivers are labeled.

It is possible different rate-limiting processes exist for different tumors and cancer types. Nevertheless, the increased burden of larger CNA or smaller INDEL deletions, and in rare cases possibly the copy number gains, are likely the early drivers. In addition, our theoretical model converged on the observed SIRs of some subgroups with fewer required drivers when estimates were based on smaller *vs.* larger deletions (*i.e.* breast cancer in *BRCA1* carriers). Thus, even within deletions, specific mutational processes may play a larger role than others depending on the status of *BRCA1* and *BRCA2* and tissue context^10^. The exact mechanisms of *BRCA1* and *BRCA2* deficiency are likely diverse.

### Comparison to genomic drivers prioritized using other methods

Existing methods mostly estimate drivers from either theoretical calculations or observed sequencing data, and few studies directly compared estimates of total driver counts derived from these methods^40^. To offer an observational comparison, we derived the average driver gene SNV and CNA profile for 1,070 breast and 566 ovarian cancer patients in TCGA using two driver prioritization methods described in Bailey *et al.*^31^ and Sanchez-Vega *et al.*^32^. Briefly, both methods focused on focal changes in known cancer driver genes. The prioritized CNAs considered only recurrent amplifications or deep (possibility homozygous) deletions and were further required to correlate with mRNA abundance changes.

Figure 4b-c shows the genes with the highest average SNV and CNA driver count per patient. The average total SNV driver count was higher for both breast cancer (Figure 4b) and ovarian cancer (Figure 4c) using Bailey *et al.*’s prioritization method^31^ compared to Sanchez-Vega *et al.*’s prioritization method^32^ (n_breast_: 2.43 *vs.* 1.13; n_ovarian_: 1.80 *vs.* 1.35; Figure 4b-c). As expected, both driver prioritization methods identify, on average, a minimum of one to three SNV drivers per patient. Both gene prioritization methods revealed *TP53* (Freq_Bailey_ *_et al._*: 0.36; Freq_Sanchez-Vega_ *_et al._*: 0.35; Figure 4b) and *PIK3CA* (Freq_Bailey_ *_et al._*: 0.37; Freq_Sanchez-Vega_ *_et al._*: 0.34; Figure 4b) to be the two most commonly mutated driver genes amongst breast cancer patients, while mutations in *TP53* (Freq_Bailey_ *_et al._*: 0.92; Freq_Sanchez-Vega_ *_et al._*: 0.95; Figure 4c) amongst ovarian cancer patients were far more common than in any other gene. *TP53* mutations are highly associated with aneuploidy, which often involves substantial genomic deletions^41^.

The aforementioned driver prioritization methods predominantly focused on point mutations and focal events. We additionally compared our driver requirement estimates to the reconstructed evolutionary histories of 198 breast and 113 ovarian tumors from the ICGC-TCGA Pan Cancer Analysis of Whole Genomes (PCAWG)^33^. Gerstung *et al.* quantified the relative timing of 28 and 26 recurrent somatic events (*i.e.* those occurring in >5% of samples) in breast and ovarian cancer, respectively. By determining the likelihoods of relative ordering between pairs of somatic mutations, they identified mutational events that preferentially occur early or late during tumor evolution. Comparing the relative timing of these somatic mutations to their recurrence rate, deletions were the most recurrent with a subset occurring preferentially early during tumor evolution.

Considering somatic mutations in the earliest quartile, 4/7 and 5/7 were deletions in breast and ovarian cancer, respectively (Figure 4d-e). Deletions of chromosomes 13q (location of *RB1* and *BRCA2*) and 17 (location of *BRCA1*) – 17p for *BRCA –* were highly recurrent, early events in both cancer types. In ovarian cancer, 4q and 22q deletions were early events that each affected over 80% of cases. The deletion of 16q occurs early and affects more than 60% of breast cancer patients. In comparison, while SNVs and indels in *PIK3CA* and *CBFB* were predicted to be amongst the earliest occurring mutations in breast cancer, they were observed in only 24.4% and 6.1% of the samples, respectively (Figure 4d). They may be drivers that are only required and frequently observed for specific subtypes of breast tumors (*i.e.*, *PIK3CA* mutation in luminal breast tumors). SNVs and indels in *TP53* were amongst the most recurrent and earliest occurring mutations in both breast and ovarian cancer (recurrence_breast_ = 47.5%; recurrence_ovarian_ = 92.0%; Figure 4d-e). *TP53* mutations are associated with increased aneuploidy and are likely facilitating the deletion phenotype. The orthogonal mutation timing data support deletions as a critical rate-limiting mutation process in breast and ovarian cancer initiation.

Finally, we sought to identify genomic overlaps between early deletions identified through the evolution-timing analysis and driver deletions prioritized by Sanchez-Vega *et al.*^32^ (**Supplementary Table 2**). We note that Sanchez-Vega *et al.* strictly considered events that were statistically recurrent, correlated with mRNA abundance, and showed a GISTIC value of -2 indicating deep (possibly homozygous) deletions, and thus may have an elevated false-negative rate. In breast cancer, the most recurrent, early deletion is chromosome 17p, in which *TP53* is located. In ovarian cancer, the predominant chromosome 17 deletion may additionally be associated with the prioritized *NF1* (located on 17q) deletion found in 7.9% of cases. *RB1*, whose prioritized deletion is found in 4.1% of breast tumors and 9.0% of ovarian tumors, is located on 13q—detected as another early, recurrent deletion by evolutionary analyses in both cancer types. Prioritized *PTEN* deletion affects 5.1% of breast tumors and intersects with the early 10q23.31 deletion. Some recurrent chromosome-arm deletions do not overlap with prioritized deletions of gene-level drivers and may affect other genes or trigger systematic effects. Overall, these analyses highlight selected chromosomal deletions (*i.e.*, 17 and 13q) and their intersections with cancer driver genes in breast and ovarian tumorigenesis. Thus, multiple independent approaches converge on a subset of deletions that may trigger tumorigenesis in breast and ovarian cancer. Implicating deletions as the rate-limiting trigger suggests somatic point mutations in cancer genes observed in normal tissue may be less relevant in tumor initiation^12,13^. This has important consequences for early detection of breast and ovarian cancers.

### Evaluation of Rate-limiting Genomic Alterations at a Cellular Level

Having characterized the event rates in large patient cohorts, we next examined the rates of SNVs, deletions, and amplifications using single-cell datasets of breast and ovarian cancer/pre-cancer cells. First, we analyzed a recently generated single-cell whole-genome-sequencing (scWGS) dataset of several CRISPR-engineered 184-hTERT L9 cell lines: SA1054 (*BRCA1-/-*), SA1055 (*BRCA2-/-b*), SA1056 (*BRCA2*-/-a), SA1292 (*BRCA1+/-*), SA1188 (*BRCA2+/-*), SA906a (*TP53-/-a*), SA906b (*TP53-/-b*), and SA039 (WT), each with at least 377 cells. All cells were harvested between passages 20-59 and verified for their genotypes, enabling us to conduct comparative analysis of SNV and CNV burdens and measure the effect of homozygous/heterozygous loss of *BRCA1/BRCA2/TP53* (**Figure 5a, Methods**).

**Figure 5:**
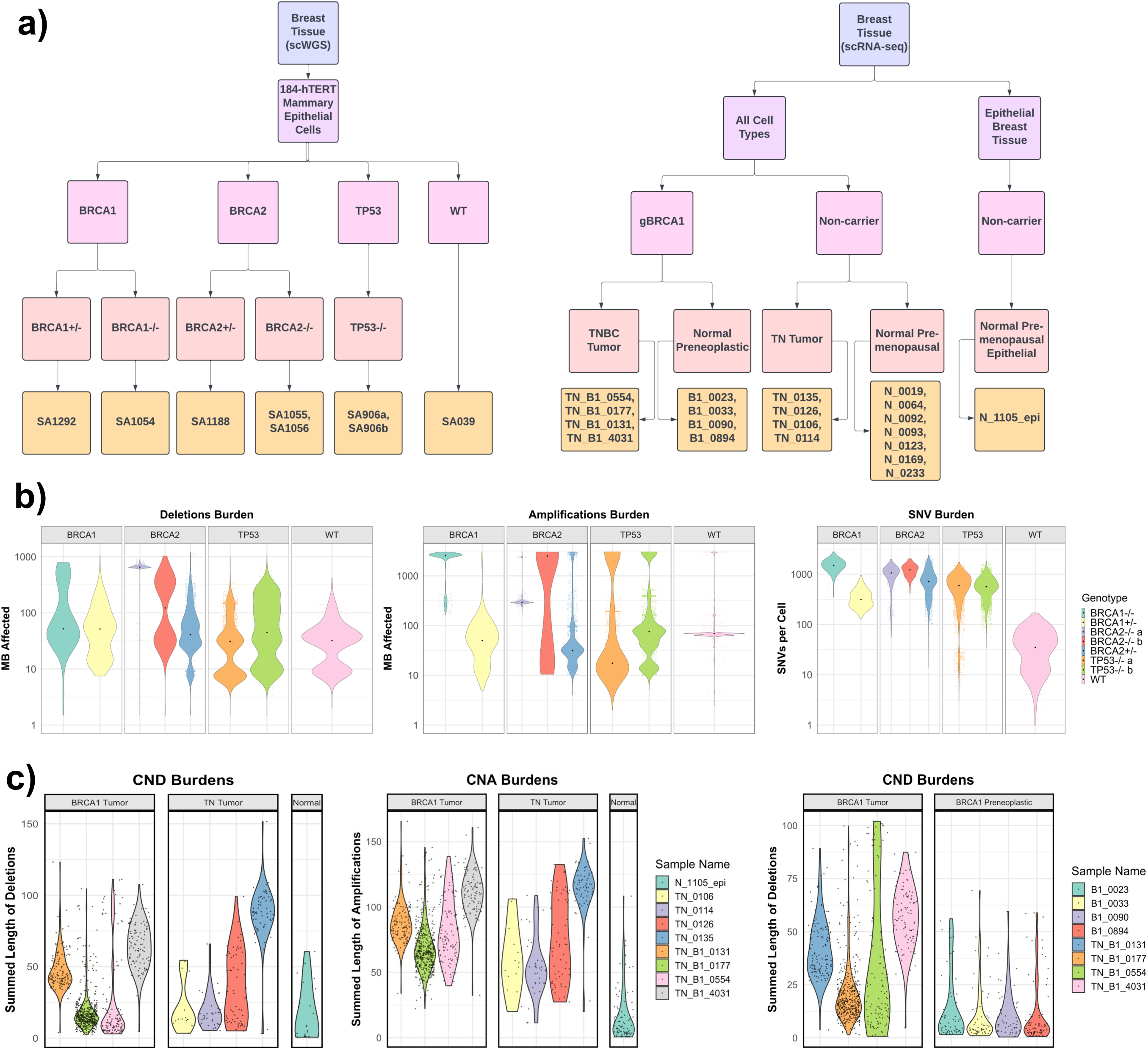
Single-cell comparison of rate-limiting mutational process in gBRCA1/2 vs non-carrier tumor samples and genetically engineered cells. a) Summary flowchart of single cell datasets and samples used for analyses, including in the left panel, data from Funnell, T., et al. (2022) that provided scWGS-analyzed 184-hTERT Mammary Epithelial Cells with genetically-engineered *BRCA1, BRCA2*, and *TP53* genotypes. On the right panel, Pal, B. et al. (2021) provided scRNA-Seq-analyzed samples from gBRCA1 carriers and non-carriers, including Triple Negative Breast Cancer (TNBC) tumor tissues, premenopausal/pre-neoplastic normal breast tissues. b) Cell-level comparison of deletion, amplification, and SNV burdens in genetically-engineered *BRCA1, BRCA2, TP53*, and WT cells included in the scWGS dataset. c) Cell-level comparison of deletion and amplification burdens of TNBCs from gBRCA1 carriers and non-carriers (left) and gBRCA1 TNBCs and pre-neoplastic normal breast tissues included in the scRNA-Seq dataset, where CNAs were inferred using inferCNV. For (b-c), each point provides the genomic alteration burden of a given cell in the sample. For amplifications/deletions, the CNA burdens were calculated by the summed length of megabase affected. The dot in each violin represents the median cell in a given sample.

For SNV burdens, all CRISPR-engineered cells exhibited significantly higher loads compared to WT cells (Wilcoxon Rank Sum test, FDR < 0.01)(Figure 5b). *BRCA1*-/- cells exhibited a significant increase in SNVs with a median fold change of 42.8X, *BRCA2*-/-a cells at 30.4X, and *BRCA2*-/-b cells at 35X. Heterozygous *BRCA1*+/- cells showed 8.93X, *BRCA2*+/- at 20.5X, whereas *TP53*-/-a cells and *TP53*-/-b are at 17.0X and 16.2X, respectively. For the burden of amplifications, *BRCA1*-/- cells exhibited a significant increase with a 37.2X fold change compared to WT, *BRCA2*-/-a cells at 4.26X, and *BRCA2*-/-a cells at 36.43X. Heterozygous *BRCA1*+/- and *BRCA2*+/- cells both demonstrated a decrease in amplifications at 0.72X (FDR < 0.01) and 0.46X (FDR < 0.05), respectively. *TP53*-/-a and *TP53*-/-b cells showed 0.25X and 1.09X amplification burdens compared to WT, respectively. For the burden of deletions, *BRCA1*-/- cells showed a significant increase with a 1.61X fold change compared to WT cells, *BRCA2*-/-a cells at 20.36X, and *BRCA2*-/-b cells at 3.83X (FDR < 0.01). Heterozygous *BRCA1*+/- and *BRCA2*+/- cells exhibited a significant but lesser increase in deletions at 1.60X and 1.27X fold change (FDR < 0.05), respectively. On the other hand, *TP53*-/-a cells and *TP53*-/-b cells showed 0.96X and 1.40X deletions of the WT. Overall, *BRCA1*/2-engineered cells showed increased SNV burdens similar to *TP53*-/- cells, but uniquely exhibited increased burdens of amplifications (in only homozygous cells) and deletions (in both heterozygous and homozygous cells) compared to WT cells.

We next analyzed triple-negative breast cancer (TNBC) tissue samples analyzed in the same scWGS study, which included cells from 4 FBI (enriched in fold-back inversions), 2 HRD-Dup (enriched in small tandem duplications), and 1 TD (enriched in large tandem duplications) TNBC samples (**Supplementary Figure 7a**). The HRD-Dup group included one germline *BRCA1* sample labeled g*BRCA1* and a somatic *BRCA1* sample labeled s*BRCA1*. For deletions, cells from the g*BRCA1* TNBC sample had significantly more deletions compared to cells from the 4 FBI TNBCs (median fold change 4.53X, Wilcoxon Rank Sum test, FDR < 0.01) and the 1 TD TNBC (2.34X, FDR < 0.01). On the other hand, g*BRCA1* cells had fewer amplifications compared to the FBI group (0.3X, FDR < 0.01) and the TD subgroup (0.41X, FDR < 0.01). For SNVs, the g*BRCA1* TNBC cells had 1.15X SNVs (FDR = 0.097) compared to the FBI group and fewer SNVs compared to the TD subgroup, with a median fold change of 0.85 (FDR = 0.014) (**Supplementary Figure 7b**).

The High-Grade Serous Ovarian Cancer (HGSC) dataset in the same scWGS study contained cells from 8 FBI samples, 6 HRD-Dup samples, and 2 TD samples (**Supplementary Figure 7a**). The HRD-Dup group included one g*BRCA1* sample and one s*BRCA1* sample. For deletions, cells from other samples of the HRD-Dup group exhibited significantly more deletions compared to the g*BRCA1* sample, with a fold change of 7.88 (FDR < 0.01). For SNVs, cells from the HRD-Dup group and other groups exhibited fewer SNVs compared to the g*BRCA1* sample (FDR < 0.01). Amplification events showed less variability across HRD statuses, with other samples from the HRD-Dup group having fewer amplifications compared to the g*BRCA1* sample, with a fold change of 0.205 (FDR < 0.01) (**Supplementary Figure 7c**).

We also analyzed a single-cell RNA-Seq (scRNA-Seq) dataset that includes *gBRCA1* carrier (N=4) vs. non-carrier (N=4) TNBC tumor samples, as well normal breast tissue samples from preneoplastic *BRCA1^+/-^* carriers (N=4) and premenopausal non-carriers (N=8)(Figure 5a). One normal premenopausal epithelial breast tissue sample in the same dataset (N_1105_epi) was used as a reference for InferCNV to obtain somatic CNVs from scRNA-Seq data (**Methods**). Compared to other TNBCs, *gBRCA1* TNBC cells exhibited 9.12X deletions (FDR < 0.01) and 11.7X amplifications (FDR < 0.01) in median fold change (Figure 5c). CNV landscapes at the cellular levels were consistent within the four *gBRCA1* TNBC samples, whereas one non-carrier TNBC sample (TN_0135) was an outlier with increased CNV (**Supplementary Figure 8**). There may also exist minor differences in CNA burdens between *gBRCA1* carriers and noncarriers in normal tissue samples, albeit less pronounced: g*BRCA1* preneoplastic cells showed 1.35X amplifications (FDR < 0.01) and 1.17X deletions (FDR = 0.97) when compared to non-carrier normal premenopausal samples (**Supplementary Figure 9**).

Finally, we compared tumor and normal samples within the same genotype to assess genomic changes throughout tumorigenesis. In g*BRCA1* carriers, g*BRCA1* TNBC cells showed 8.06X deletions and 3.90X amplifications (FDR < 0.001) compared to g*BRCA1* preneoplastic cells (Figure 5c). In comparison, non-carrier TNBCs had 1.39X deletions (FDR < 0.01) and less amplifications at 0.47X (FDR < 0.01) when compared to non-carrier premenopausal cells (**Supplementary Figure 9**). Overall, these data confirm at a cellular level that g*BRCA1/2* TNBCs and engineered cells typically showed higher CNA burdens than non-carrier or WT cells, and deletions are the most consistent type of mutation process that show higher rates in g*BRCA1/2* cells.

## DISCUSSION

Cancer is caused by the successive accumulation of aberrations that deregulate normal cellular processes and trigger tumorigenesis^1,2^. However, despite multiple efforts, the driving mutational process and the number of aberrations required to initiate a diagnosable tumor remain largely unknown^3–11^. Identifying the trigger of tumorigenesis would help delineating malignant from non-malignant somatic evolution and provide key information for accurate risk stratification of pre-malignant lesions. Integrating established theoretical and empirical methods, we compared mutational processes in BRCA-heritable breast and ovarian tumors to their sporadic counterparts. We focused on breast and ovarian cancer to avoid confounding with well-known sex-biases in cancer mutations and to ensure sufficient numbers of carriers for modeling (n > 10)^42,43^. Our data suggested deletions as the rate-limiting mutational process initiating tumorigenesis in breast and ovarian cancer. Despite current protocols largely focused on point mutations, these data prioritize structural variants and copy number alterations in early detection strategies, such as liquid biopsies.

Although prioritized drivers based on sequencing studies (Figure 4) include many classes of alterations (*e.g. PIK3CA* hotspots, *ERBB2* amplifications)^31,32^, these may not represent rate-limiting steps required to become a diagnosable breast or ovarian tumor. Similarly, profiling somatic mutations in non-malignant tissues have demonstrated a high prevalence of mutations in cancer genes, *e.g. NOTCH*, despite the absence of cancer^12,13^. While these mutations may confer a fitness advantage, they are unlikely to be the rate-limiting events in early tumorigenesis. Reconstructing the history of breast and ovarian tumors predicted multiple highly recurrent deletions as early events many of which were arm-level deletions that intersected with tumor suppressor genes, including *RB1* on 13q as well as *NF1* and *TP53* on chromosome 17. Interestingly, a recent study of cancer/non-cancer breast tissues identified a key clonal event to be derivative chromosome der(1;16), which includes the entire long arm of chromosome 1 (1q) and a truncated short arm of chromosome 16 (16p), acquired during early puberty to late adolescence in breast cancer patients. *TP53* mutational status is also known to be correlated with aneuploidy^41^. However, reconstructing the history of breast and ovarian tumors predicted more early deletions than our theoretical framework predicted. Because timing estimates are rank-based, we cannot estimate the time-frame between the acquisition of two drivers. Despite both being labeled early events, one deletion event could occur long before a second early event. Further, it is possible not all deletions confer a fitness advantage.

*BRCA1* or *BRCA2* loss of function leads to HRD and higher dependence on more error prone repair pathways, such as non-homologous end joining (NHEJ) and microhomology mediated-end joining (MMEJ). HRD tumors are enriched in <50 kbp genomic deletions characteristic of NHEJ or MMEJ repair^10,39^. Specifically, recent ICGC PCAWG studies identified an indel signature ID6 that correlated with the SBS3—often attributed to defective HRD; ID6 is characterized predominantly by deletions of ≥5 base pairs and with overlapping microhomology at the boundaries, suggesting a potential deletion type that may occur more frequently when tumors develop HRD deficiency.

The mechanisms by which *BRCA1* and *BRCA2* deficiency promotes tumorigenesis are likely myriad. The penetrance of *BRCA1* and *BRCA2* deficiency can be modulated by the genomic environment^44^, and their role in tumorigenesis may be compounded by the tumor microenvironment^45^, both of which are not fully captured in our framework. Our framework assumes both subgroups require a similar number of driver alterations and does not model the scenario where carriers may require one fewer driver alteration than non-carriers. We focus on determining the lower-bound on the rate-limiting number of exonic driver alterations required. Future work may also consider genomic rearrangements, non-coding, transcriptional or post-transcriptional changes that could contribute to tumorigenesis. We could not differentiate clonal from subclonal mutations, with the former likely being more informative of the rate-limiting requirements for tumorigenesis^33^.

*BRCA1* and *BRCA2* carriers are recommended to begin screening for breast cancer starting at age 25 and discussions of risk-reducing mastectomy or salpingo-oophorectomy starting at age 35^27^. There is an increased prevalence of pre-malignant lesions, *e.g.* ductal carcinoma in-situ (DCIS) and tubual intraepithelial carcinoma (TIC), in *BRCA1* and *BRCA2* carriers^28–30^. With this increased prevalence comes a need to accurately identify pre-malignant lesions at high-risk of transformation into invasive disease. Understanding the rate-limiting processes triggering transformation could be directly leveraged for risk-stratification and early detection.

## METHODS

### Data

#### TCGA *BRCA1* and *BRCA2* status in breast and ovarian cancer

Patients with pathogenic variants in *BRCA1* and *BRCA2* were identified by the TCGA PanCanAtlas Germline project^35^. To avoid confounding of somatic differences by ancestry, we only considered individuals of European descent.

#### TCGA sample somatic mutation rate

We downloaded the public MAF somatic mutation call file from the TCGA PanCanAtlas MC3 project^34^ that conducted using a variety of variant calling tools. We then filtered the file to retain only non-synonymous mutations (in one of these types: “Missense_Mutation”, “Nonsense_Mutation”, “Splice_Site”, “Translation_Start_Site”). The mutation rate is calculated as the number of non-synonymous mutations divided by the age at the initial diagnosis for each sample.

#### TCGA sample copy-number variation rate

We downloaded the ABSOLUTE-annotated seg file from the TCGA PanCanAtlas Aneuploidy project^41^ (https://gdc.cancer.gov/about-data/publications/pancan-aneuploidy). CNA burden was calculated as both the number of segments and the number of base-pairs involved in a CNA. Both measures were normalized by the age of initial diagnosis for each sample. Segments and altered bases were further categorized as being involved in a gain or a deletion event.

#### TCGA sample small INDEL mutation rate

We downloaded the public MAF somatic mutation call file from the TCGA PanCanAtlas MC3 project^34^ that conducted using a variety of variant calling tools. We then filtered the file to retain only small insertions and deletions. The INDEL burden is calculated as both the total number of INDELs and the total number of bps in an insertion or deletion divided by the age at the initial diagnosis for each sample.

#### Standardized incidence ratios for breast and ovarian cancer

We performed an extensive literature search for standardized incidence ratios (SIRs) for both breast and ovarian cancer amongst carriers of *BRCA1* and *BRCA2*. We utilized SIR estimates derived by Kuchenbaecker *et al.* for a variety of reasons^38^. The multinational cohort used to calculate these SIRs was the largest amongst all studies reviewed, the range of ages represented was greater than most, and the study was one of the most recent [**Supplementary Table 1**]. The estimates listed below are total SIRs through age 80.

Breast Cancer:

> Total SIR (95% CI) *BRCA1*: 16.6 (14.7-18.7)
>
> Total SIR (95% CI) *BRCA2*: 12.9 (11.1-15.1)

Ovarian Cancer:

> Total SIR (95% CI) *BRCA1*: 49.6 (40.0-61.5)
>
> Total SIR (95% CI) *BRCA2*: 13.7 (9.1-20.7)

#### Incidence ratio estimation

We estimated the incidence ratio using a framework proposed by Tomasetti *et al*.^6^. Briefly, Tomasetti *et al.* showed there is a power relationship between cancer incidence and the number of rate-limiting mutational events for the initiation of a cancer subgroup and the average mutation rate of the cancer subgroup^6^. For example, a cancer subgroup that has a mutation rate twice that of another subgroup will have an incidence rate 2^n^ that of the second subgroup, where n represents the number of rate-limiting mutational events required for tumorigenesis. Here we consider *BRCA1* and *BRCA2* carriers compared to sporadic breast and ovarian cancers. We randomly sampled a subset of non-carrier breast cancer patients to match the sample size of *BRCA1* carriers with breast cancer. Non-carrier breast cancer patients were also matched to the clinical characteristics of the carrier cohort, *i.e.* we ensured the same proportion of patients of each stage, grade and subtype in both the carrier and non-carrier groups. Next, we calculated the ratio between the median non-carrier mutation rate and the median *BRCA1* carrier mutation rate. Finally, we estimated the incidence ratio considering the mutation rate ratio, u, and various number of driver events, k, using the following:

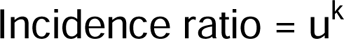

Because each additional driver event is likely to add a growth advantage and assuming this growth advantage is constant with each successive driver, as done in Tomasetti *et al*.^6^ based on the modeling of self-renewal phase of a tissue^46^ and tumor clonal expansion^47^, the incidence ratio prediction becomes:

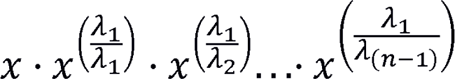

Where: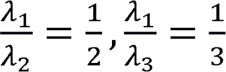, *etc*

We repeated these randomly sampling 10,000 times and compared the median across the 10,000 iterations to the observed SIRs. We considered the driver number that minimized the Kolmogorov–Smirnov metric between the estimated and observed SIR distribution to be the required number of drivers. We concluded a model failed to converge if the Kolmogorov– Smirnov metric never reached a minimum after testing 15 drivers. We repeated the same process with ovarian cancer patients. We considered mutation rates based on SNVs, CNAs, CNA deletions, CNA gains and INDEL deletions. For CNAs and INDELs, we calculate the total number of bps involved in these aberrations.

#### Mutational timing analysis

We leveraged mutational timing estimates from Gerstung *et al.*^33^. Briefly, for 28 and 26 recurrent somatic events (defined as occurring in >5% of samples) in breast and ovarian cancer, respectively, Gerstung *et al.* generated a discrete multinomial distribution based on pairwise probabilities one mutation occurs before another mutation, (x_i_, x_j_) ∼ p1, p2, p3.

p1 = P(x_i_ before x_j_)

p2 = P(x_j_ before x_i_)

p3 = P(order of (x_i_,x_j_) unknown)

Using league modeling, they stimulated games by drawing from this multinomial distribution and assigning points based on “wins”, *i.e.* mutation A comes before mutation B, or “ties”, *i.e.* the order of mutations unknown. The final scores were ranked to generate relative timing estimates, and this process was repeated at least 1,000 times to obtain distributions of the rankings. Here, these relative timing estimates were compared to the recurrence of the mutation across samples of the same cancer type within the PCAWG cohort.

#### CNV frequency maps

To generate the CNA frequency maps of *BRCA1*, *BRCA2*, and non-carrier tumors, TCGA SNP array data was downloaded from the GDC repository (https://dcc.icgc.org/releases) genome version GRCh38. Data category ‘Masked Copy Number Segment’ derived from SNP6.0 genotyping arrays. CNA frequency maps are generated using the R package svpluscnv (https://github.com/ccbiolab/svpluscnv) with the following options cnv.freq(x, fc.pct=0.2), where ‘x’ is the object containing segmentation data and ‘fc.pct’ is the percentage of CNA log-R ratio change threshold defining gain and loss of copy number. i.e CNA gain (red) threshold is logR > log2(1+ fc.pct) and loss (blue) is logR < log2(1-0.2). The same measures were used to determine chromosome arm CNA frequencies reported samples with at least 30% of the chromosome arm length gained or lost.

### Single-cell whole-genome sequencing (scWGS) dataset and analysis

We analyzed three distinct datasets from Funnell, T., et al. that conducted single-cell Whole Genome Sequencing (scWGS) on genetically engineered Mammary Epithelial Cells, Triple Negative Breast Cancer (TNBC), and High-Grade Serous Ovarian Cancer (HGSC) datasets. The 184-hTERT Mammary Epithelial Cells datasets included clonal populations with genetically engineered *BRCA1*, *BRCA2*, and *TP53* deletions produced via CRISPR–Cas9 editing of 184-hTERT L9 cell lines: SA1054 (*BRCA1-/-*) with 382 cells, SA1055 (*BRCA2-/-b*) with 391 cells, SA1056 (*BRCA2*-/-a) with 496 cells, SA1292 (*BRCA1+/-*) with 377 cells, SA1188 (*BRCA2+/-*) with 472 cells, SA906a (*TP53-/-a*) with 650 cells, SA906b (*TP53-/-b*) with 984 cells, and SA039 (WT) with 878 cells. The TNBC dataset includes 4 FBI-dup samples, 2 HRD samples, and 1 TD sample. The HGSC dataset contained 8 FBI-dup samples, 6 HRD samples and 2 TD samples.

Based on the scWGS data, CNV segment sizes were calculated in base pairs and converted to megabases (MB) affected by copy number deletions and amplifications, respectively, and subsequently the median fold changes for these CNVs and SNVs across different groups were calculated using R. The Wilcoxon Rank Sum test was employed to statistically discern differences in CNV and SNV distributions across groups or phenotypes. P values were adjusted for multiple testing using the BH procedure for FDR.

### Single-cell RNA-seq (scRNA-Seq) dataset and analysis

Seurat objects of single-cell RNA-seq data were obtained from human breast tissue from 21 patients with no history of cancer treatment as previously described. This scRNA-Seq dataset was generated with the 10x Genomics Chromium platform and an illumina NextSeq sequencer. The samples were taken from patients at different stages of cancer progression, including normal healthy cell samples as well as various tumor samples.

The paper provided data on triple-negative breast cancer (TNBC) tissues of germline *BRCA1* carriers (g*BRCA1*) vs. non-carriers, as well as preneoplastic and normal breast tissue. TNBC taken from four *BRCA1* carriers were labeled “B1”. The other TNBCs were labeled “TN”. The seurat object containing both samples, *SeuratObject_TNBC.rds*, was further divided into four *BRCA1* variant samples and four non-*BRCA1* variant samples. The 1 normal premenopausal epithelial breast tissue sample (N_1105_epi) used as a reference sample was from the *SeuratObject_NormEpi.rds* object. Data from precancerous and pathologically normal breast tissue was sourced from the seurat object *SeuratObject_NormB1Total.rds*. Lastly, the four samples of preneoplastic normal breast tissue that was taken from four *BRCA1* variant carriers were labeled “PN” and the non-carrier premenopausal normal breast tissue samples were labeled “TP”.

Using the R package, inferCNV, we inferred CNVs of *BRCA1*-carrier TNBCs, non-carrier TNBCs, and their respective normal counterparts. The inferCNV package analyzes the intensity of gene expression across the genome of a tumor sample and compares these values to a designated normal somatic cell sample as a reference. The i6 Hidden Markov Model (HMM) was implemented in the inferCNV process to assign six states of copy number variation to each tumor cell. The states 2 and 1 were used in the creation of violin plots for deletion as they correspond to a loss of one and two copies, respectively. The same was done with amplification as states 4 and 5 represent the gain of one and two copies. CNV segment sizes were calculated in base pairs and converted to megabases (MB) affected by copy number deletions and amplifications, respectively, and subsequently the median fold changes for these CNVs across different groups were calculated. The Wilcoxon Rank Sum test was employed to statistically discern differences between groups, and P values were adjusted for multiple testing using the BH procedure for FDR.

#### Data visualization

Visualizations were generated in the R statistical environment (v3.3.1) with the lattice (v0.24-30), latticeExtra (v0.6-28) and BPG (v5.6.23) packages^48^.

#### Data availability

Processed mutation calls were downloaded from https://gdc.cancer.gov/about-data/publications/pancanatlas. To generate CNA frequency maps, data was downloaded from the GDC repository (https://dcc.icgc.org/releases).

#### Code availability

Code to generate results can be found: https://github.com/khoulahan/brca-driver-estimation

## Supplementary Figure Legends

**Supplementary Figure 1: Consistent tumor purity and ploidy across subgroups**

**a)** The number of samples that experienced genome doubling was consistent across subgroups. The density plot shows the distribution of sample-wise genome doubling for each subgroup. **b)** Tumor purity was not significantly different between cancer subgroups with the exception of *BRCA2* ovarian cancer samples were had significantly lower tumor purity than wildtype. Density plot shows the distribution of sample-wise tumor purity for each subgroup.

**Supplementary Figure 2: Increased mutation rates in *BRCA1* and *BRCA2* carriers**

Breast **(A)** and ovarian **(B)** tumors in *BRCA1* and *BRCA2* carriers show increased mutation counts and rates, defined as count/age, compared to non-carrier (WT) individuals. Boxplots represent median, 0.25 and 0.75 quantiles with whiskers are 1.5x interquartile range. P-values from the Kruskal-Wallis test. GAIN: gains; INSERT: insertions

**Supplementary Figure 3: Mutation burden ratio distributions**

Distributions of mutation burden ratios between carrier and non-carrier over all of the 10,000 bootstrap iterations for SNVs **(a)**, CNAs **(b)**, CNA deletions **(c)**, small deletions **(d)** and gains **(e)**.

**Supplementary Figure 4: Estimating incidence from CNA segments**

Incidence estimates based on CNA segment counts **(a-d)**, CNA deletion segment counts **(e-h),** INDEL deletion counts **(i-l)** and CNA gains segment counts **(m-p)**. All segment counts are normalized by age at diagnosis.

**Supplementary Figure 5: Estimating breast cancer incidence in the METABRIC cohort** Estimated incidence rates based on SNV and CNA (gain and losses) in the METABRIC breast cancer cohort.

**Supplementary Figure 6: Copy number frequency map of *BRCA1*, *BRCA2*, and non-carrier tumors**

Genome-wide representation of CNVs indicating the number of samples (left axis) and frequencies (right axis) with copy number loss (light blue/blue) and gain (salmon/red). The lighter colors represent short chromosome arm (p) and intense color represents long (q) arms. **a-c)** Breast cancers carriers for *BRCA1* **(a)**, *BRCA2* **(b)** and wild type **(c)**. **d-f)** Ovarian cancer samples carriers for *BRCA1* **(d)**, *BRCA2* **(e)** and wild type **(f)**.

**Supplementary Figure 7: Single-cell comparison of rate-limiting mutation process in triple negative breast cancer and high-grade serous ovarian cancers.** a) Summay flowchart of scWGS datasets from Funnell, T. et al. (2022). TNBC and HGSC cells with gBRCA1, sBRCA1 and other miscellaneous genotypes are outlined here. b) Cell-level comparison of deletion, amplification and SNV burden in gBRCA1, sBRCA1 and miscellaneous genotypes of both TNBC and c) HGSC cells in the scWGS dataset. Each dot represents the either the CNV burden of an individual cell in terms of metabases affected or the number of SNV events per cell (b-c).

**Supplementary Figure 8: inferCNV-generated heatmaps of CNV in both TNBC and normal samples of gBRCA1 carriers and non-carriers based on scRNA-Seq data.** Heat maps of CNV landscapes generated by the R package inferCNV, including that for g*BRCA1* TNBCs (a), non-carrier TNBCs (b), g*BRCA1* preneoplastic normal tissue (c) and non-carrier premenopausal normal samples (d). Amplifications for each chromosome region are highlighted red, while deletions are highlighted blue. The normal reference sample is at the top of each plot, while each sample analyzed is below (a-d).

**Supplementary Figure 9. Single-cell comparison of rate-limiting mutational process in non-carrier TNBC, non-carrier premenopausal and gBRCA1 preneoplastic samples based on inferCNV analyses of scRNA-Seq data.** Cell-level comparison of non-carrier tumor and normal samples in terms of CND (a) and CNA (b) burdens. CND (c) and CNA (d) burdens compared across gBRCA1 preneoplastic and non-carrier premenopausal cells. b) Individual dots of the violin plots represent the summed length of megabases deleted or amplified for a given cell (a-d).

## Supplementary Tables Legends

**Supplementary Table 1: Standardize incidence rate (SIR) of breast and ovarian cancer in BRCA1 and BRCA2 carriers**

**Supplementary Table 2: Early CNA deletions identified through evolution-timing analysis and driver prioritization methods**

Overlap between early deletions identified through the evolution-timing analysis and driver deletions prioritized by Sanchez-Vega *et al.*^32^ AVG_CNA_PER_PT = average CNA per patient; AMP = amplification; DEL = deletion

**Supplementary Table 3: Single cell datasets and samples used for analyses**

The list of samples is from (1) Funnell, T. et al. (2022) that provided scWGS data of genetically engineered 184-hTERT Mammary Epithelial Cells, TNBC, and HGSC samples of various genotypes; (2) Pal, B. et al. (2021) that provided scRNA-Seq-analyzed samples from gBRCA1 carriers and non-carriers, including Triple Negative Breast Cancer (TNBC) tumor tissues and premenopausal/pre-neoplastic normal breast tissues.

## ACKNOWLEDGEMENTS

K.E.H. was supported by a CIHR Vanier Fellowship. This work was supported by the NIH/NCI under award number P30CA016042 and by an operating grant from the National Cancer Institute Early Detection Research Network (1U01CA214194-01). This work was also supported by NIH NIGMS R35GM138113 and ACS RSG-22-115-01-DMC to K.H. P.V.L. is a CPRIT Scholar in Cancer Research and acknowledges CPRIT grant support (RR210006). P.V.L. was supported by the Francis Crick Institute, which receives its core funding from Cancer Research UK (CC2008), the UK Medical Research Council (CC2008), and the Wellcome Trust (CC2008). The authors wish to acknowledge patients and families participating in The Cancer Genome Atlas and the International Cancer Genome Consortium (ICGC). They also thank the TCGA PanCanAtlas, PCAWG working groups, and Drs. Marc Williams and Sohrab Shah for providing their studies’ data and ensuring meaningful use.

## CONTRIBUTIONS

Performed statistical and bioinformatics analyses: K.E.H., M.B., J.G.C., D.F., G.L.G., H.H.

Data Processing: K.E.H., D.F., M.W., S.H.

Wrote the first draft of the manuscript: K.E.H., D.F., K.H.

Edited the manuscript: K.E.H., P.V.L., P.C.B., K.H.

Initiated the project: K.E.H., K.H.

Supervised research: P.C.B., K.H.

Approved the manuscript: all authors

## REFERENCES

1. B. Vogelstein, N. Papadopoulos, V. E. Velculescu, S. Zhou, L. A. Diaz Jr, K. W. Kinzler, Cancer genome landscapes. Science 339, 1546–1558 (2013).

2. L. A. Garraway, E. S. Lander, Lessons from the cancer genome. Cell 153, 17–37 (2013).

3. W. Y. Tan, X. W. Yan, A new stochastic and state space model of human colon cancer incorporating multiple pathways. Biol Direct 5, 26 (2010).

4. M. P. Little, Are two mutations sufficient to cause cancer? Some generalizations of the two-mutation model of carcinogenesis of Moolgavkar, Venzon, and Knudson, and of the multistage model of Armitage and Doll. Biometrics 51, 1278–1291 (1995).

5. S. Wu, W. Zhu, P. Thompson, Y. A. Hannun, Evaluating intrinsic and non-intrinsic cancer risk factors. Nature Communications 9, (2018).

6. C. Tomasetti, L. Marchionni, M. A. Nowak, G. Parmigiani, B. Vogelstein, Only three driver gene mutations are required for the development of lung and colorectal cancers. P Natl Acad Sci USA 112, 118–123 (2015).

7. M. H. Bailey, et al. Comprehensive Characterization of Cancer Driver Genes and Mutations. Cell 173, 371–385 (2018).

8. I. Martincorena, et al. Universal Patterns of Selection in Cancer and Somatic Tissues. Cell 171, 1029–1041 (2017).

9. S. Saghafinia, M. Mina, N. Riggi, D. Hanahan, G. Ciriello, Pan-Cancer Landscape of Aberrant DNA Methylation across Human Tumors. Cell Rep 25, 1066–1080, (2018).

10. Y. Li, et al. Patterns of somatic structural variation in human cancer genomes. Nature 578, 112–121 (2020).

11. G. Ciriello, et al. Emerging landscape of oncogenic signatures across human cancers. Nature Genetics 45, 1127–1133 (2013).

12. I. Martincorena, et al. Somatic mutant clones colonize the human esophagus with age. Science 362, 911–917 (2018).

13. H. Lee-Six, et al. The landscape of somatic mutation in normal colorectal epithelial cells. Nature 574, 532–537 (2019).

14. C.O. Nordling, A new theory on cancer-inducing mechanism. Br J Cancer 7, 68–72 (1953).

15. P. Armitage, R. Doll, The age distribution of cancer and a multi-stage theory of carcinogenesis. Br J Cancer 8, 1–12 (1054).

16. A.G. Knudson Jr, Mutation and cancer: statistical study of retinoblastoma. P Natl Acad Sci USA 68, 820–823 (1971).

17. D.E. Rushlow, et al. Characterisation of retinoblastoma without RB1 mutations: genomic, gene expression, and clinical studies. Lancet Oncol 14, 327–334 (2013).

18. E. Levy-Lahad, E. Friedman. Cancer risks among *BRCA1* and *BRCA2* mutation carriers. Br J Cancer 96, 11–15 (2007).

19. R. Roy, J. Chun, S. N. Powell, *BRCA1* and *BRCA2*: different roles in a common pathway of genome protection. Nat Rev Cancer 12, 68–78 (2011).

20. P. Jonsson et al. tumor lineage shapes BRCA-mediated phenotypes. Nature 571, 576–579 (2019).

21. A. Lal, D. Ramazzotti, Z. Weng, K. Liu, J. M. Ford, A. Sidow, Comprehensive genomic characterization of breast tumors with *BRCA1* and *BRCA2* mutations. BMC Med Genomics 12, 84 (2019).

22. S. Nik-Zainal, et al. Landscape of somatic mutations in 560 breast cancer whole-genome sequences. Nature 534, 47–54 (2016).

23. M. M. Kamieniak, et al. DNA copy number profiling reveals extensive genomic loss in hereditary *BRCA1* and *BRCA2* ovarian carcinomas. Br J Cancer 108, 1732–1742 (2013).

24. D. Yang, S. Khan, Y. Sun, K. Hess, I. Shmulevich, A. K. Sood, W. Zhang, Association of *BRCA1* and *BRCA2* mutations with survival, chemotherapy sensitivity, and gene mutator phenotype in patients with ovarian cancer. JAMA 306, 1557–1565 (2011).

25. R.A. Taylor, et al. Germline *BRCA2* mutations driver prostate cancers with distinct evolutionary trajectories. Nat Commun 8, 13671 (2017).

26. N. Turner, A. Tutt, A. Ashworth. Hallmarks of ‘BRCAness’ in sporadic cancers. Nature Reviews Cancer 4, 814–819 (2014).

27. National Comprehensive Cancer Network. Genetic/Familial High-Risk Assessment: Breast and Ovarian (Version 3.2019). (2019). Retrieved from: https://www2.tri-kobe.org/nccn/guideline/gynecological/english/genetic_familial.pdf

28. Hoogerbrugge, N., et al. High prevalence of premalignant lesions in prophylactically removed breasts from women at hereditary risk of breast cancer. J. Clin. Oncol. 21, 41–45 (2003).

29. Hermsen, B.B.J., et al. Low prevalence of (pre) malignant lesions in the breast and high prevalence in the ovary and Fallopian tube in women at hereditary high risk of breast and ovarian cancer. Int. J. Cancer. 119, 1412–1418 (2006)

30. Mingels, M.J.J.M., et al. Tubal epithelial lesions in salpingo-oophorectomy specimens of BRCA-mutation carriers and controls. Gynecol. Oncol. 127, 88–93 (2012).

31. M. H. Bailey, et al. Comprehensive characterization of cancer driver genes and mutations. Cell 173, 371–385 (2018).

32. F. Sanchez-Vega, et al. Oncogenic signaling pathways in The Cancer Genome Atlas. Cell 173, 321–337 (2018).

33. M. Gerstung, et al. The evolutionary history of 2,658 cancers. Nature 578, 122–128 (2020).

34. K. Ellrott, et al. Scalable open science approach for mutation calling of tumor exomes using mulitiple genomic pipelines. Cell Syst. 6, 271–281 (2018).

35. K. L. Huang, et al. Pathogenic germline variants in 10,389 adult cancers. Cell 173, 355–370 (2018).

36. J. Yuan, et al. Integrated analysis of genetic ancestry and genomic alterations across cancers. Cancer Cell 34, 549–560 (2018).

37. L.A. Forsberg et al. Age-related somatic structural changes in the nuclear genome of human blood cells. Am J Hum Genet 90, 217–228 (2012).

38. K. B. Kuchenbaecker, et al. Risks of breast, ovarian and contralateral breast cancer for *BRCA1* and *BRCA2* mutation carriers. JAMA 317, 2402–2416 (2017).

39. H. Davies, et al. HRDetect is a predictor of *BRCA1* and *BRCA2* deficiency based on mutational signatures. Nat Med 23, 517–525 (2017).

40. C. J. Tokheim, N. Papadopoulos, K. W. Kinzler, B. Vogelstein, R. Karchin, Evaluating the evolution of cancer driver genes. P Natl Acad Sci USA 113, 14330–14335 (2016).

41. A. M. Taylor, et al. Genomic and functional approaches to understanding cancer aneuploidy. Cancer Cell 33, 676–689 (2018).

42. Y. Yuan, et al. Comprehensive characterization of molecular differences in cancer between male and female patients. Cancer Cell 29, 711–722 (2016).

43. C.H. Li et al. Sex differences in cancer driver genes and biomarkers. Cancer Res 78, 5527–5537 (2018).

44. A. C. Fahed, et al. Polygenic background modifies penetrance of monogenic variants conferring risk for coronary artery disease, breast cancer, or colorectal cancer. 10.1101/19013086 (29 November 2019).

45. A. A. Kraya, et al. Genomic signatures predict the immunogenicity of BRCA-deficient breast cancer. Clin Cancer Res 25, 4363–4374 (2019).

46. C. Tomasetti, B. Vogelstein, G. Parmigiani, Half or more of the somatic mutations in cancers of self-renewing tissues originate prior to tumor initiation. P Natl Acad Sci USA 110, 1999–2004 (2013).

47. R. Durrett, S. Moseley, Evolution of resistance and progression to disease during clonal expansion of cancer. Theor Popul Biol 77, 42–48 (2010).

48. C. P’ng, et al. BPG: seamless, automated and interactive visualization of scientific data. BMC Bioinformatics 20, 42 (2019).

