## Supplementary Figures for "Deletions Rate-Limit Breast and Ovarian Cancer Initiation"

Supplementary Figure 1

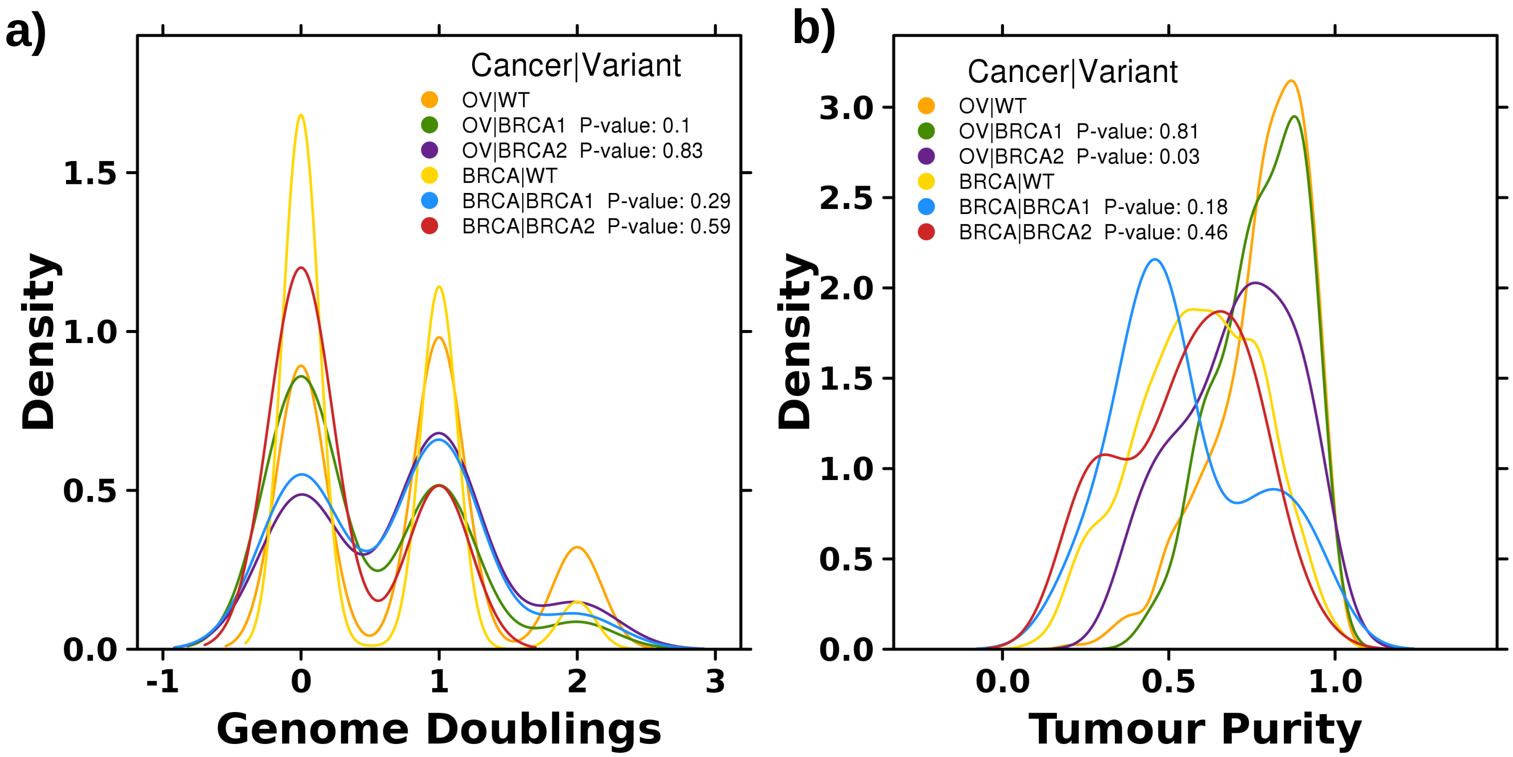

### Supplementary Figure 2

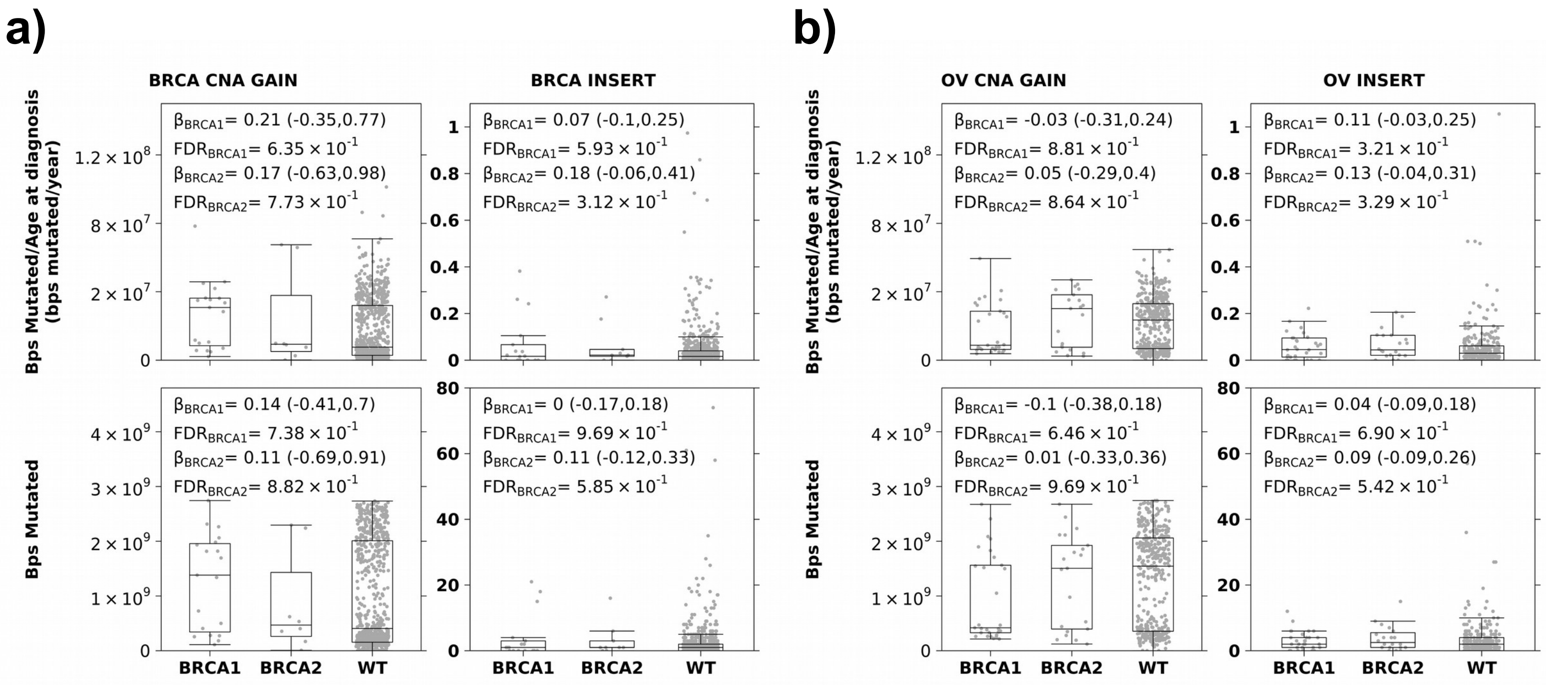

Supplementary Figure 3

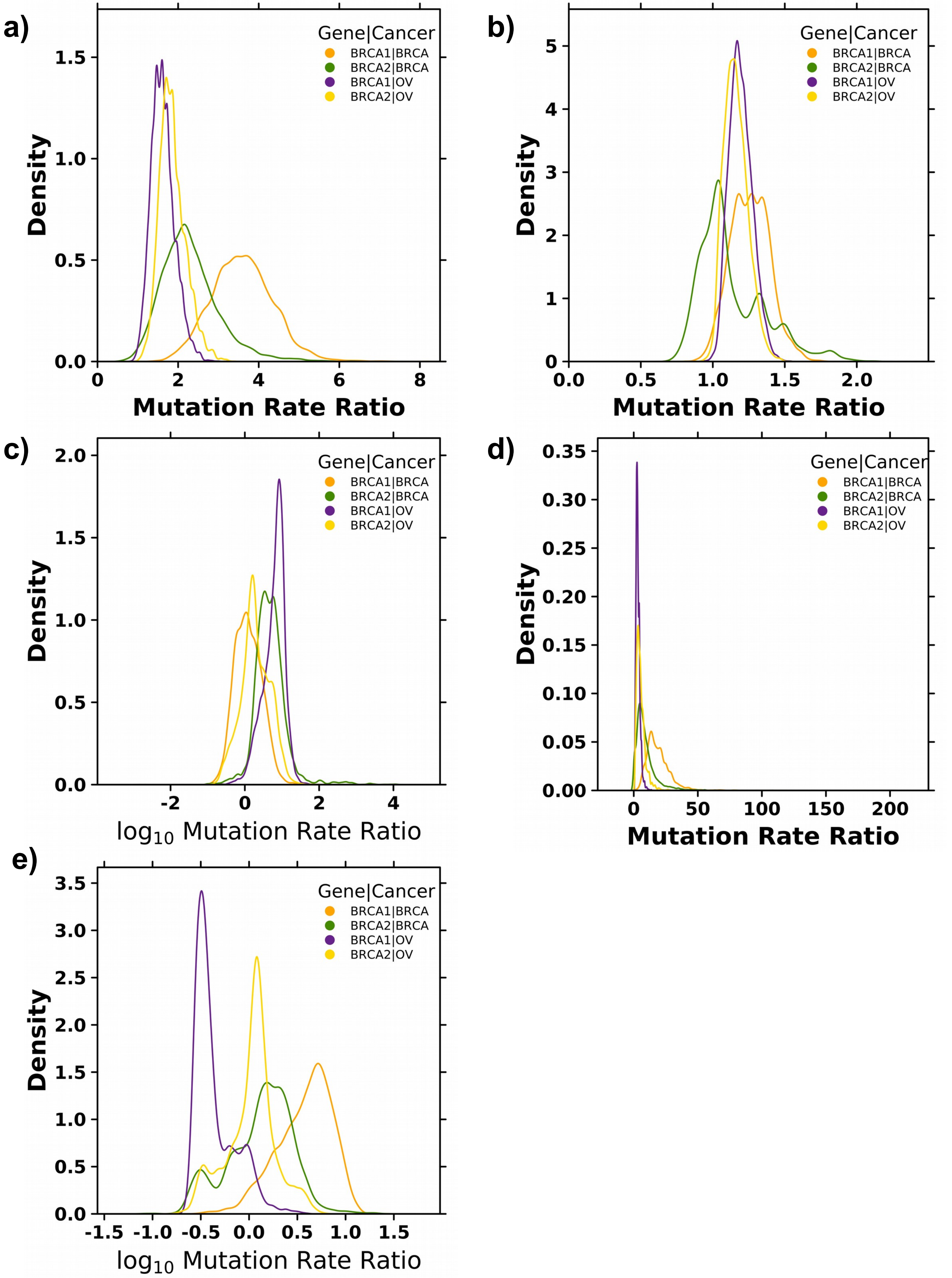

Supplementary Figure 4

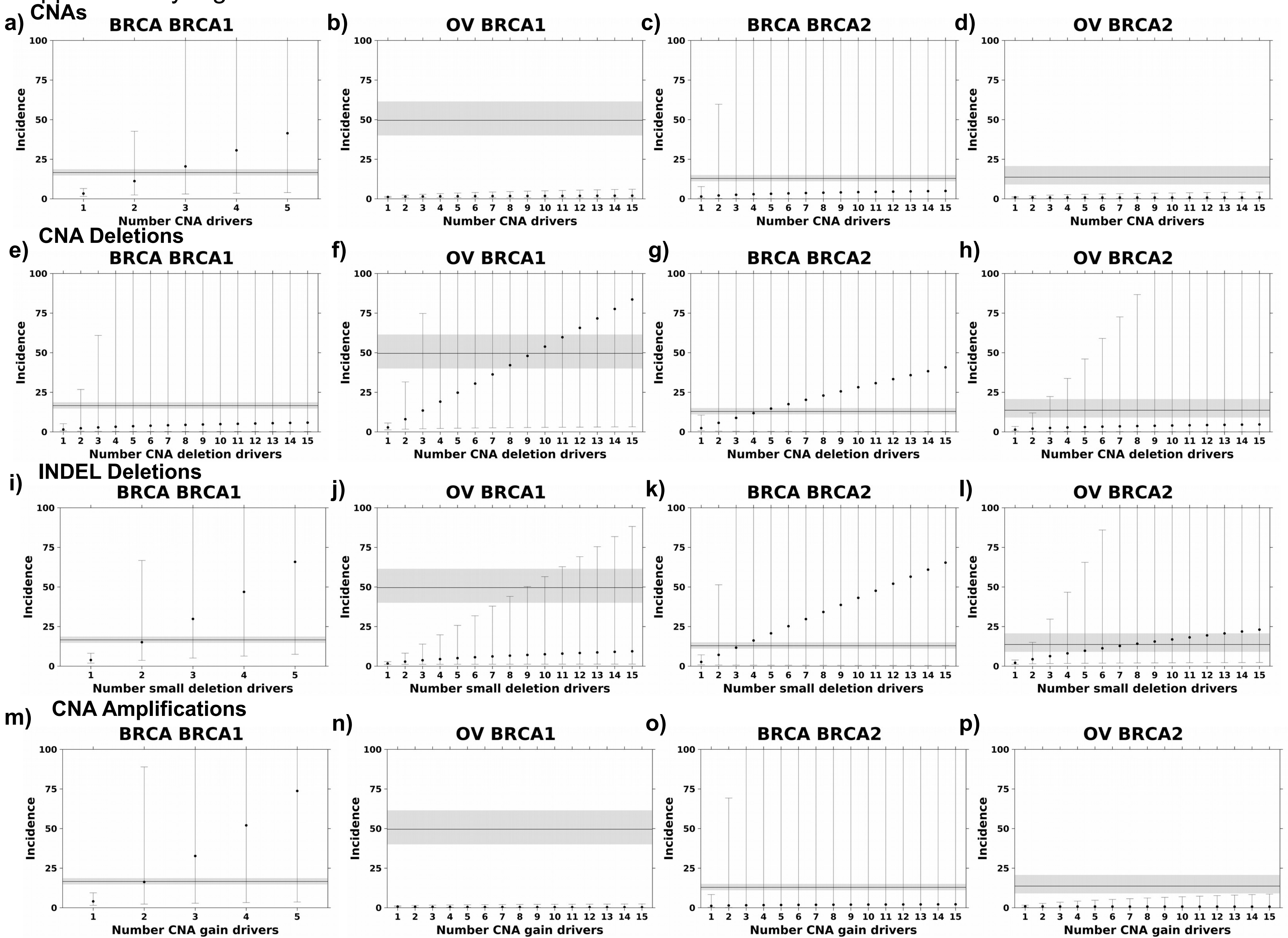

### Supplementary Figure 5

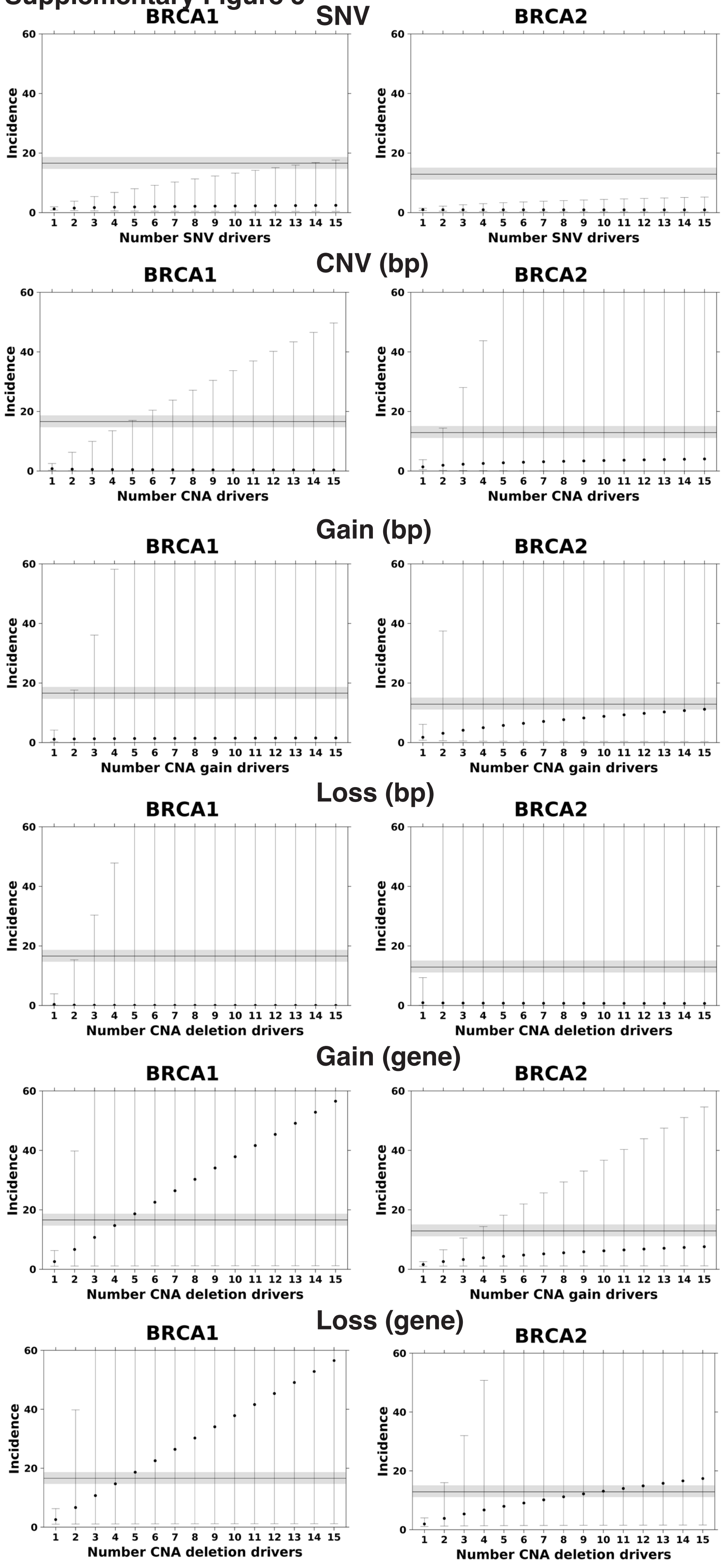

Supplementary Figure 6

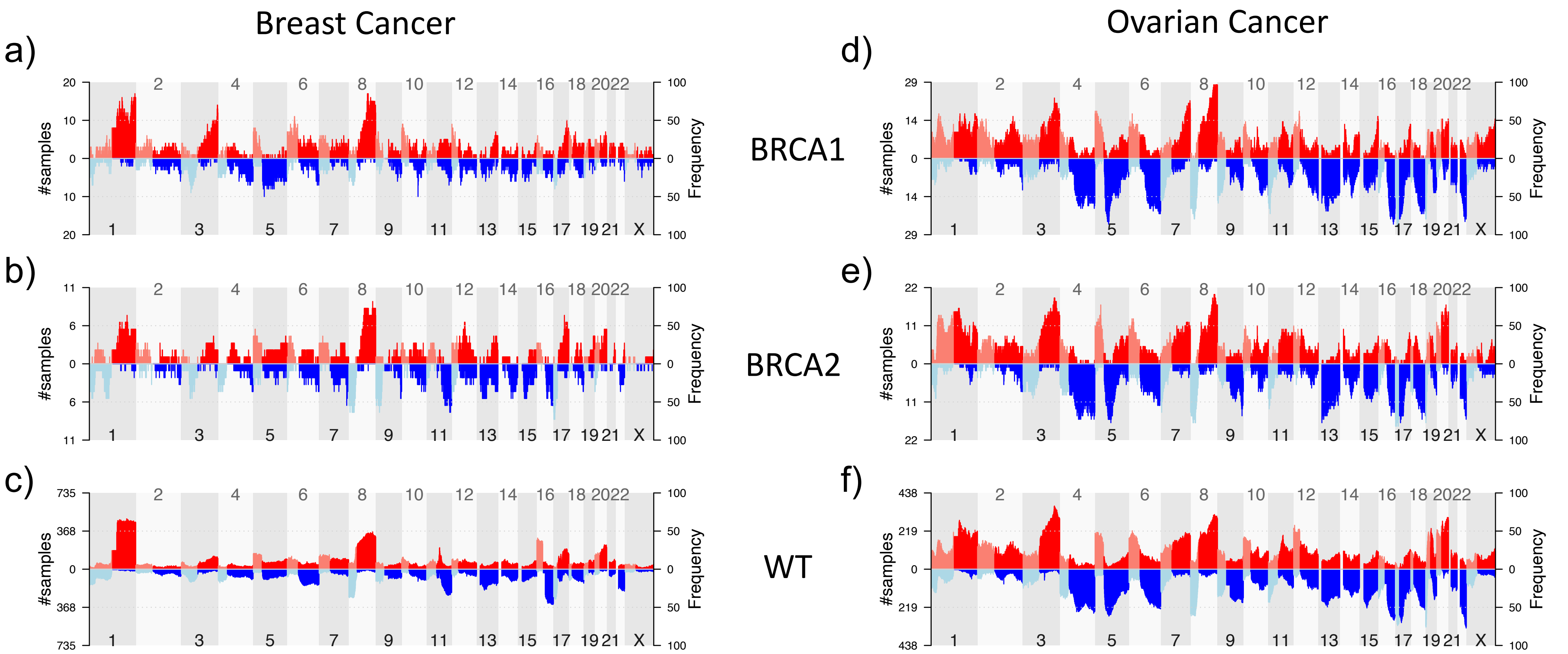

### Supplementary figure 7

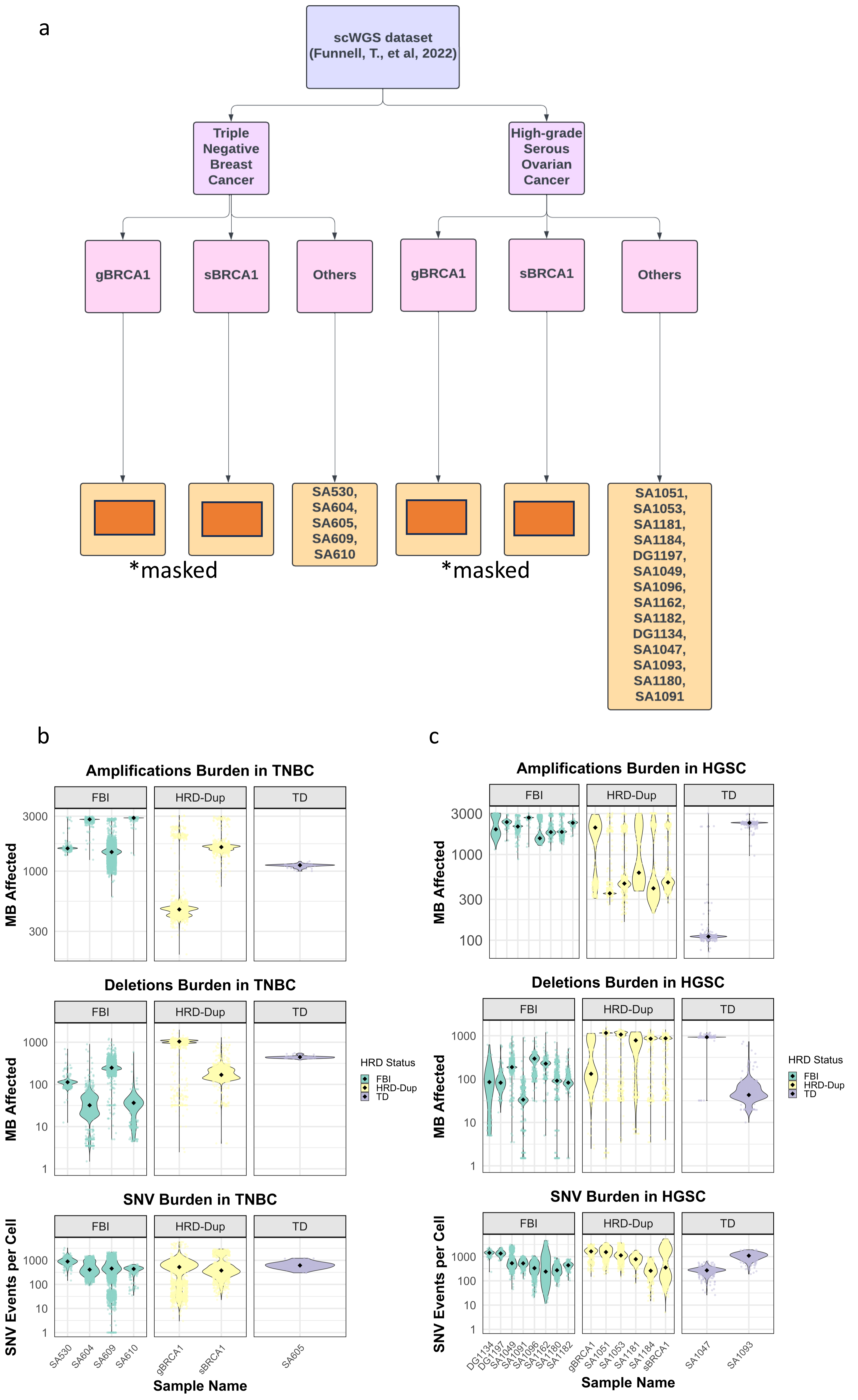

### Supplementary figure 8

a. inferCNV heatmap of BRCA1 TNBC Samples (Normal Epithelial Reference)

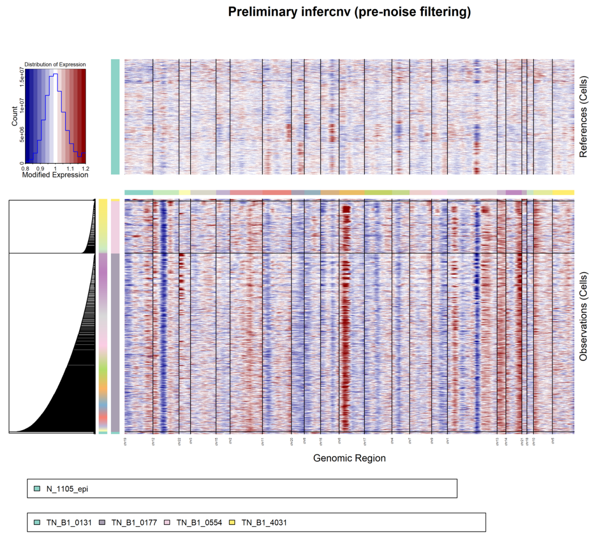


b. inferCNV heatmap of non-BRCA1 TNBC Samples (Normal Epithelial Reference)

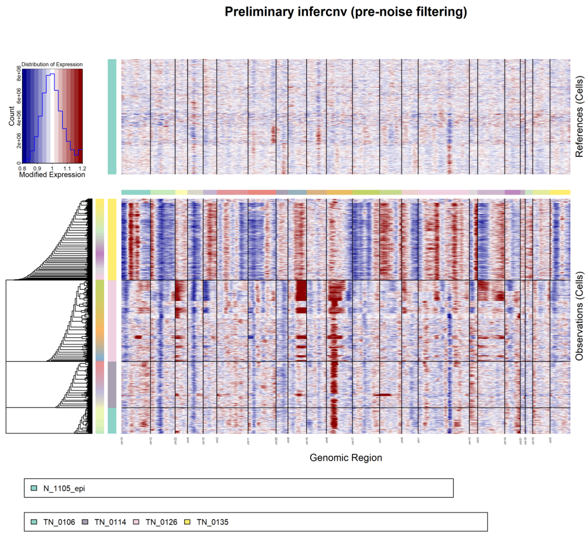


c. inferCNV heatmap of BRCA1+/- Preneoplastic Normal Breast Tissue (Normal Epithelial Reference)

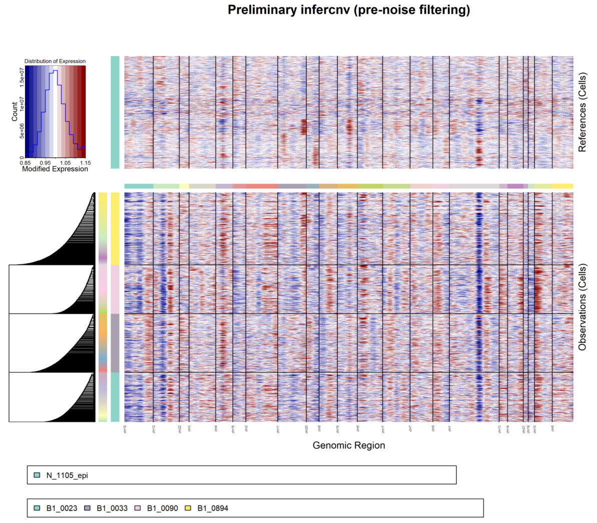


d. inferCNV heatmap of non-BRCA1 Premenopausal Normal Breast Tissue (Normal Epithelial Reference)

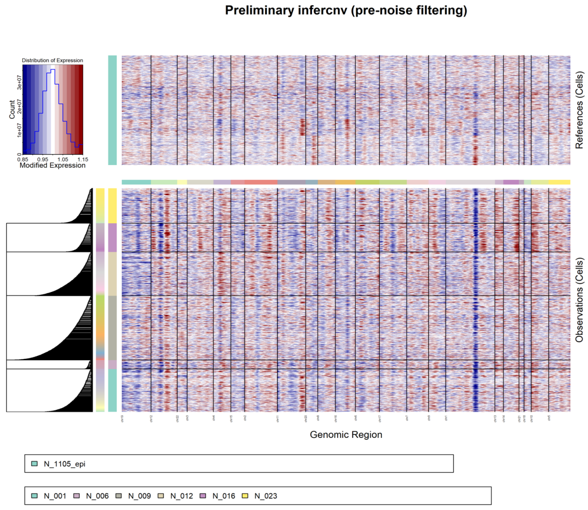

### Supplementary figure 9

a. CND Length of non-BRCA1 TNBCs & non-BRCA1 Premenopausal Normal Breast Tissue

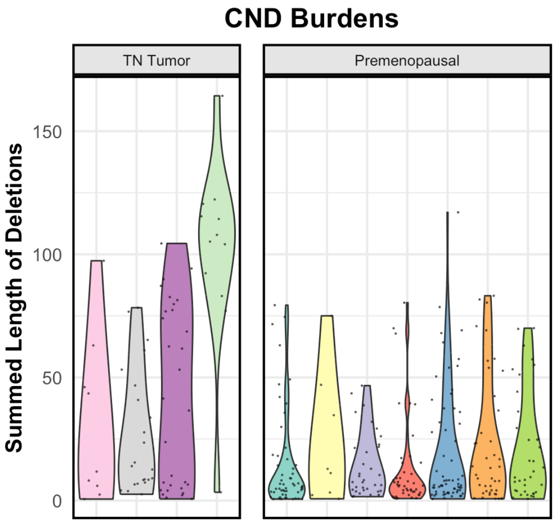


b. CNA Length of non-BRCA1 TNBCs & non-BRCA1 Premenopausal Normal Breast Tissue

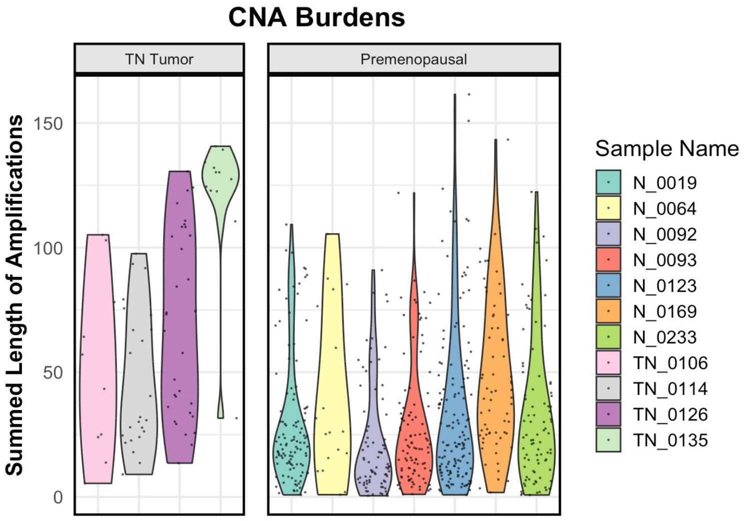


c. CND Length of BRCA1+/- Preneoplastic Normal Breast Tissue & non-BRCA1 Premenopausal Normal Breast Tissue

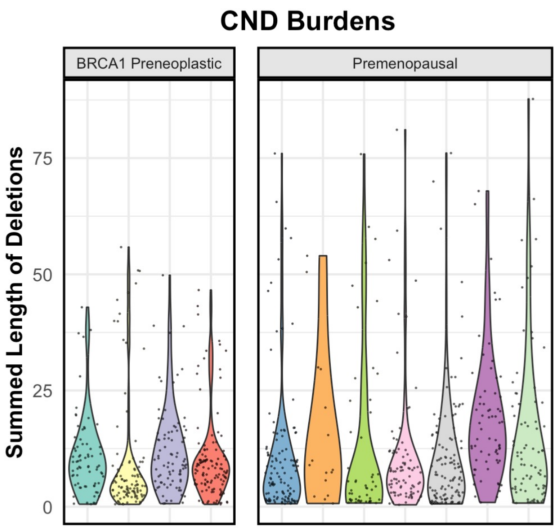


d. CNA Length of BRCA1+/- Preneoplastic Normal Breast Tissue & non-BRCA1 Premenopausal Normal Breast Tissue

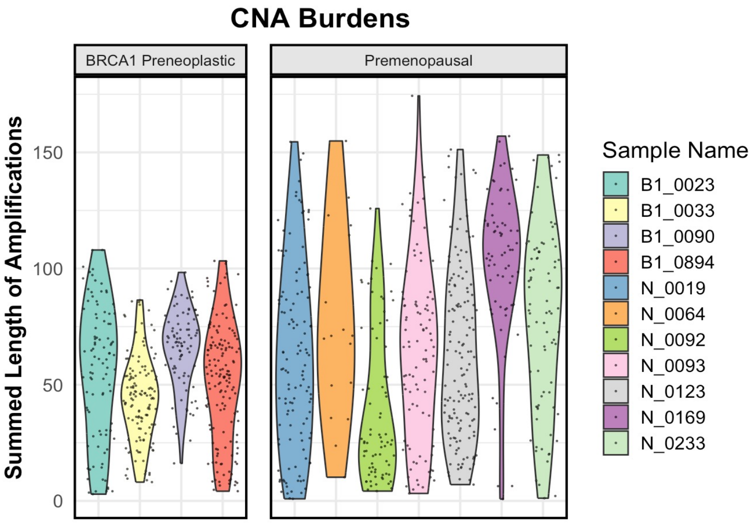
